# Glucocorticoids modulate expression of perineuronal net component genes and parvalbumin during development of mouse cortical neurons

**DOI:** 10.1101/2025.02.12.637910

**Authors:** Liang Yue, Michael T Craig, Brian J Morris

**Affiliations:** School of Psychology and Neuroscience, College of Medical, Veterinary and Life Sciences, University of Glasgow, Sir Joseph Black Building, Glasgow, G12 8QQ, UK

**Keywords:** neurocan, mifepristone, RU486, Has3, Bral2, non-genomic

## Abstract

Severe prenatal maternal stress is a risk factor for schizophrenia in offspring. Since parvalbumin-containing GABAergic interneuron function in cortex and hippocampus is compromised in schizophrenia, and perineuronal nets (PNNs) facilitate the functioning of these cells, we tested the hypothesis that glucocorticoids, as stress mediators that can access the foetal compartment, might influence the expression of PNN component genes. In cultured mouse cortical neurons, we detected effects of hydrocortisone on many PNN component genes, via diverse mechanisms. A rapid (<4h), glucocorticoid receptor (GR)-mediated suppression of *neurocan* and hyaluronan synthase (*Has*) 1 and 3 mRNAs was observed at 7 days *in vitro* (DIV), whereas at 14DIV, *brevican* and *versican* expression was reduced by hydrocortisone without GR involvement, while GR inhibition elevated *Has1* and *Has2* mRNA levels and suppressed *aggrecan* mRNA levels. *Tenascin R* expression was rapidly suppressed by hydrocortisone at 7DIV but not at 14. At 21DIV, PNN component gene expression had become insensitive to hydrocortisone, although *parvalbumin* expression was reduced after 24h but not 4h exposure. Additionally, effects on protein levels were observed that were sometimes consistent with the mRNA changes (e.g. Has3, Gad1) and sometimes unrelated to them (e.g. elevated TnR levels at 14DIV after glucocorticoid receptor antagonism). We found that hydrocortisone could directly inhibit proteasome activity, potentially explaining the ability of hydrocortisone to increase Has2 levels. As expected from these results, the overall structure of the PNN was compromised by hydrocortisone exposure, with the length of proximal dendrite covered by PNN being reduced. Overall, the data demonstrate a complex and profound, but developmental stage-dependent, regulation of PNN component gene expression by glucocorticoids. This may contribute to the action of severe prenatal or perinatal stress to increase schizophrenia risk.

## Introduction

Perineuronal nets (PNNs) are complex structures composed of extracellular matrix molecules, covering the somata, dendrites and proximal axon segments of distinct neuronal populations (Fawcett et al., 2019), mainly enveloping fast-spiking GABAergic parvalbumin (*Pvalb*)-expressing interneurons in cortical and hippocampal areas (Celio & Chiquet-Ehrismann, 1993). Their formation in the developing brain is associated with the onset of *Pvalb* expression, the maturation of the *Pvalb*-expressing cells, and the closure of the critical period of cortical plasticity (Morishita et al., 2015; Pizzorusso et al., 2002; Willis et al., 2022). Once formed, the PNNs are believed to enable fast-spiking activity of these cells by restricting ion movements immediately extracellular to the cell membrane (Fawcett et al., 2019; Morawski et al., 2015), while also protecting the cells from oxidative stress (Morishita et al., 2015) and consolidating afferent synapse architecture (Ferrer-Ferrer & Dityatev, 2018).

The key components of PNNs are the members of the family of chondroitin sulphate proteoglycans (CSPGs), including aggrecan (*Acan*), brevican (*Bcan*), neurocan (*Ncan*), versican (*Vcan*) and phosphacan (*Ptprz1*), which are essential for PNN function and structure (Deepa et al., 2006). Hyaluronan (HA), which forms the backbone of PNNs, binding to the CSPGs, is synthesised by cell membrane hyaluronan synthase (Has) enzymes, and extruded into the extracellular space. The structure is stabilised by hyaluronan and proteoglycan link proteins (Haplns), and by cross-linking by Tenascin R (Fawcett et al., 2019). The cellular origins of these different components is not always clear, but it is likely that some are synthesised in neurons and others in astrocytes (Irala et al., 2024; John et al., 2006), suggesting that a complex regulatory framework must be required for effective coordination of the synthesis of the different constituent proteins in different cells.

During development, PNNs are susceptible to external stressors. Reduced PNN staining intensity and density were found in various brain areas, including prefrontal cortex (PFC), basolateral amygdala (Allgäuer et al., 2023; Santiago et al., 2018) and hippocampus (Riga et al., 2017) in rodents after exposure to early life stress. Consistent evidence has also demonstrated downregulated levels of CSPG molecules after exposure to stress (Koskinen et al., 2020; Li et al., 2024).

PNNs are disrupted as part of the dysfunction of *Pvalb*-expressing neurons in schizophrenia. Alongside the reduced expression of *PVALB* and the GABA synthetic enzyme *GAD1*/GAD67 in PFC, hippocampus and thalamic reticular nucleus in schizophrenia (Gonzalez-Burgos et al., 2015; Hoftman et al., 2015), PNN structure is also compromised (Enwright et al., 2016; Mauney et al., 2013; Pantazopoulos et al., 2021; Steullet et al., 2018). Maternal experience of severe stress (such as famine or natural disaster) during pregnancy increases risk of schizophrenia for the offspring *in utero* (Mawson & Morris, 2023). While the causal mechanisms are unclear, it is known that glucocorticoids (GCs), which are markedly elevated during the maternal stress response, can cross the placenta and access the foetal compartment (Fowden et al., 2022; Gitau et al., 1998; Schwartz, 1997; Wieczorek et al., 2019). Considering the reports that early life stress could influence PNN development, we considered the hypothesis that GCs might affect the expression of PNN component genes, and hence disrupt the formation or maintenance of PNNs and increase schizophrenia risk.

In humans, the main GC is cortisol, while in rodents, the main GC is corticosterone. The canonical GC mechanism involves binding to an intracellular receptor, either the GC receptor (GR) or mineralocorticoid receptor (MR), nuclear translocation of the steroid-receptor complex, and binding to specific sites, usually in gene promoter regions, resulting in repression or activation of gene expression (Joëls, 2018). These transcriptional effects are slow – a typical time-course would have an onset of altered mRNA levels 8-12h after GC exposure (De Kloet, 2004; Joëls, 2018), although occasionally more rapid effects can be observed. The altered levels would then be maintained for more than 24h after the original exposure. The non-genomic effects of GCs, which are thought to involve membrane receptor-mediated actions, not necessarily involving alterations in gene transcription, occur much more rapidly (< 1h) (Hollos et al., 2020; Hynes & Harvey, 2019; Joëls, 2018; Joëls et al., 2013). There is also evidence that some rapid, non-genomic effects of GCs do not involve the canonical GR; but other receptors such as GPR97 (*ADGRG3*) located on the cell membrane (Y. Q. Ping et al., 2021).

Thus, we sought to test the hypothesis that elevated GC levels would disrupt PNN formation in developing neuronal networks, and to uncover the underlying mechanisms..

## Methods

### Primary cortical neuronal culture

Mouse neurons were cultured as previously described (Cruise et al., 2000; Fuller et al., 2001). The brains were removed from C57BL/6 mouse embryos at E17 and washed in ice-cold Hanks Balanced Salt Solution (HBSS). The meninges were removed, the cortical tissues were isolated and transferred into clean ice-cold HBSS. After two more washes in clean ice-cold HBSS, the cortical tissues were transferred into 0.05% trypsin/EDTA (Gibco, 25300054) at 37 °C for 10 min. Following this, DMEM (10% HI horse serum, 1% Penicillin Streptomycin, 1% Glutamax) was added to inactivate the trypsin and tissues were centrifuged at 1500 rpm for 5 min. After removing the supernatant, DMEM (8ml/embryo) was added, and the mixture of DMEM and cortical tissues were transferred into clean neurobasal medium (Gibco, 21103049) with B27 supplement (Gibco, 17504044). The cells were then seeded into 12-well plates precoated with 4 μg/ml poly-D-lysine and 6 μg/ml laminin. Subsequently, neurobasal medium with B27 supplement and clean DMEM medium were added to each well. The ratio of cell suspension medium, clean neurobasal medium and DMEM medium in each well was 0.25/0.25/0.5. After 24 h, 50% medium in each well was replaced with new neurobasal medium with B27 supplement; and then 50% of medium in each well changes with Neurobasal/B27 were made every 4 days for the duration of the cultures.

### Drug treatment

To investigate the effect of glucocorticoids on PNN expression at 7, 14 and 21 days in vitro (DIV), cells were treated with low dose (final concentration 20 nM) or high dose (final concentration 100 nM) hydrocortisone acetate (CORT) for 4h or 24h. These concentrations were chosen to reproduce those achieved in the foetus during moderate or severe maternal emotional stress (see Discussion). In some experiments, the selective GR antagonist, mifepristone/RU486 (20 nM final concentration) was added 30 min prior to CORT treatment. In addition, to examine the pharmacology of GC effects, the selective MR agonist, aldosterone (100 nM final concentration), or the selective GR agonist, fluticasone (50nM final concentration), were tested. Alternatively, cells were also treated with collagen 3 (Cell Guidance Systems, 75nM final concentration), an agonist ligand for the adhesion GPCRs GPR56/97 (Olaniru et al., 2018; Zhu, Luo, et al., 2019), to investigate whether the effect of glucocorticoids was mediated through this GPCR family. To test whether mRNA stability was affected, CORT treatment occurred together with 5µg/ml Actinomycin D treatment, to inhibit the mRNA synthesis, 1.5 h, 2 h, 3 h, or 4 h prior to mRNA extraction.

### mRNA extraction, cDNA synthesis and quantitative polymerase chain reaction (qPCR)

The procedure was according to our standard methodology (Paterson et al., 2006; Willis et al., 2021). Total mRNA was isolated from the cultured cells using RNeasy mini kits (Qiagen 74106). The RNA quality and concentration were confirmed using spectrophotometry before cDNA synthesis. First strand cDNA was synthesized from mRNA using high-capacity RNA-to-cDNA kit with 10 µl RT Buffer (Applied biosystem, 4387406) and 1 µl enzyme (Applied biosystem, 4387406), in a final volume of 20 μL with appropriate volumes of Nuclease water and mRNA samples based on the results of RNA spectrophotometry. The product was aliquoted and stored at −20 °C for future use. The cDNA quality and concentration were confirmed using spectrophotometry before qPCR method.

mRNA levels were measured using SYBRgreen methodology. Samples were run in triplicate on 96-well plates with 1 μl of cDNA samples, 19 µl master mix (Agilent), with cycling conditions of 1 cycle 50 °C for 2 min, 1 cycle 95 °C for 2 min, 40 cycles for 30 s at 95 ° C and 10 s at 60 ° C, followed by a melt curve. *Gapdh* was employed as house-keeping gene to normalise the gene expression of different targeted primers, except in a single case where the treatment (collagen 3) was found to alter *Gapdh* expression, where *Tbp* was used instead. The primers targeted *Acan, Bcan, Ncan, Vcan, Ptprz1, Has1, Has2, Has3, TnR, Hapln4, Gad1, Gad2* and *Pvalb*. Data were analysed using the ΔΔCt method. Primer sequences are provided in Supplementary table 1.

### Protein extraction and western blot

Immunoblotting was performed via our standard procedures (McNair et al., 2010; O’Kane et al., 2003). The medium was removed from the wells and 1 ml ice-cold PBS (PH7.4) was added to each well for about 1 minute. Following this, 60 μl RIPA buffer (made up with 50 mM Tris-HCL, 150 mM NaCl, 1% Triton 100, 0.15 SDS, 0.5% sodium deoxycholate and 50 mL dH2O) with 1% protease inhibitor cocktail (Sigma P-8340) and 1% Sodium orthovanadate was added to the wells for 3 min. The wells were then scraped with pipette tips, the contents were transferred to 1.5 mL Eppendorf tubes and centrifuged for 10 min at 4°C, 13,000 rpm. Supernatants were collected and the protein concentration was measured using Bradford Protein Assay (Bradford, 1976). The protein samples (−1.5μ g/μl) were prepared with 4x sample buffer (NuPAGE, Novex, NP0007) and reducing reagent (NuPAGE, Novex, NP0004). Protein samples were denatured at 80 °C for 10 min, and 25 μl/lane added to SDS-PAGE in 4%-12% Bis-Tris gels (NuPAGE, Novex, NP0302BOX), followed by electrophoresis at 200 volts for 1.5 h in chilled running buffer.

Protein was then transferred to Invitrogen PVALBDF membranes (Invitrogen, LC2002) in transfer buffer at room temperature running for 1 h at 30 V. Membranes were washed twice in ddH2O and blocked in 0.5% Tween-Tris-buffered-saline (TTBS) with 3% dried milk powder (Marvel) for 30 min at room temperature. After blocking, membranes were incubated with primary antibodies at 4°C overnight in 1% TTBS milk. The following morning, membranes were washed 3 times for 10 min in Tris-buffered-saline (TBS) containing 0.05% Tween 20 (Sigma-Aldrich, T7949) and incubated in horseradish peroxidase (HRP)-conjugated anti-mouse/anti-rabbit secondary antibodies (anti-mouse concentration: 1:10,000; anti-rabbit concentration: 1:6000 with 1% dried milk in TTBS for 1.5 −2 h at room temperature. The antibodies that were used in western blots are listed in Supplementary table 2.

Membranes were then washed once with TTBS and washed twice with TBS after incubation of secondary antibodies. Membranes with targeted antibody could be detected by adding chemiluminescent HRP Substrate (Immobilon, Millipore, WBKLS0100) using equal quantities of luminol and peroxide solution. Finally, the membranes were placed into cassette and were captured using PXI4 (Syngene) with varied exposure times depending on the antibodies used.

### Lectin fluorescence

Wisteria floribunda agglutinin (WFA) lectin is widely used as a marker to visualise PNNs (Härtig et al., 1992). For immunofluorescence, cells were fixed at 14 DIV with 4% paraformaldehyde for 30 min. Cells were then permeabilised and blocked with PBS (0.3 M NaCl)/0.25%Triton X-100/10% normal goat serum (NGS) for 1h at room temperature. Following the blocking step, cells were incubated in a humidified chamber with biotinylated WFA diluted in 0.3 M PBS/0.25%Triton X-100/3% NGS at 4°C overnight. In the following day, cells were incubated in streptavidin-conjugated Rhodamine Red-X diluted in 0.3M PBS/3% NGS for 1h in the dark. After incubation in the dark, cells were washed three times in 0.3M PBS and mounted with Vectashield mounting media (Vector Laboratories, H-1200). The slides were finally covered with a coverslip.

Images were scanned using a confocal microscope (ZEISS, LSM900) with a 10x objective for counting and 20x objective for presentation. All representative images were captured as a Z-stack (15µ M in depth) using a Z step of 0.50 µM, 20X objective lens, image size 1024×1024 pixels. The images were taken using Zen blue 3.0 software and downloaded with summed intensity zen-stack projection in Zen black system. Image analyses were performed using ImageJ (Schneider et al., 2012). The PNN-covering dendrite length and intensity were measured manually in ImageJ.

### Proteasome activity

All 3 catalytic 20S proteasome activities were measured by detecting the 7-amino-4-methylcoumarin (AMC) fluorescence liberated from the synthetic proteasomal substrates: Suc-Leu-Leu-Val-Tyr-AMC (Suc-LLVY-AMC) for chymotrypsin-like activity, Boc-Leu-Ala-Ala-AMC (Boc-LAA-AMC) for trypsin-like activity and Ac-Glu-Pro-Leu-Asp-AMC (Ac-GPLD-AMC) for caspase-like activity. To recognise whether the activities were related to the proteasome, MG132 was used as an inhibitor for all 3 activities of 20S proteasome. Unconjugated AMC was diluted into appropriate concentrations to create a standard curve which allowed the fluorescence signal to be converted to units of AMC. The released fluorescence was measured every 60 s for 30 min using excitation wavelength 340 nm and emission wavelength 450 nm.

### Statistical analysis

Statistical analysis was conducted using Minitab, with ANOVA as the standard approach after checking for normality of data distribution, with Tukey or Fisher post-hoc tests depending on the F value of the factor in ANOVA. Sample sizes given represent the number of individual culture wells from which mRNA or protein was extracted separately. Samples were nested within the culture from which they derived for statistical analysis.

## Results

Efficiency of amplification was tested for all primer pairs, and was in every case between 90 and 120% (Supplementary Information Figure S1). In addition, preliminary experiments determined that the expression of *Gapdh* mRNA was not affected by the treatments, consistent with previous studies in the laboratory (Willis et al., 2021), and was therefore an appropriate choice for “housekeeping gene” in these studies, apart from collagen 3 exposure, where *Gapdh* mRNA levels were affected by the treatment but *Tbp* mRNA levels did not change, and so was used instead (data not shown).

### CORT treatment altered the mRNA expression of PNN components during neuronal development period

The present study aims to investigate the effect of glucocorticoids on the gene expression of PNN components in mouse cortical neurones. The components tested in the study included CSPGs (*Acan*, *Bcan*, *Ncan* and *Vcan*), HAS (*Has1*, *Has2* and *Has3*), *Hapln4* (also known as Brain link protein 2/Bral2) and tenascin R (*TnR*). Attempts were also made to detect *Hapln* 1, *Hapln2* and *Hapln3*, but they appeared to be below the threshold for reliable detection (Ct’s ∼ 33 or higher).

We then studied the effect of exposure at 7, 14 and 21 DIV to low (20nM) or high dose (100nM) CORT and where effects of CORT were observed, we sought to investigate the involvement of GR activation in the actions: mifepristone (100nM) was cotreated CORT. Mifepristone has nanomolar affinity for glucocorticoid receptors, but micromolar affinity for mineralocorticoid receptors (Testas et al.,1983; Galanuad et al., 1984; Cain et al., 2019).

The mRNA expression of *Vcan* was not significantly changed by CORT exposure at 7 DIV (4h: F (2, 43) =1.02, p=0.370; 24h: F (2, 42) =0.42, p=0.66) (Fig.1 A, D), or 21 DIV (4h: F (2, 19) =3.00, p=0.085; 24h: F (2, 18) =0.11, p=0.894) after 4h and 24h CORT treatment (Fig.1 C,F).

**Figure 1.**
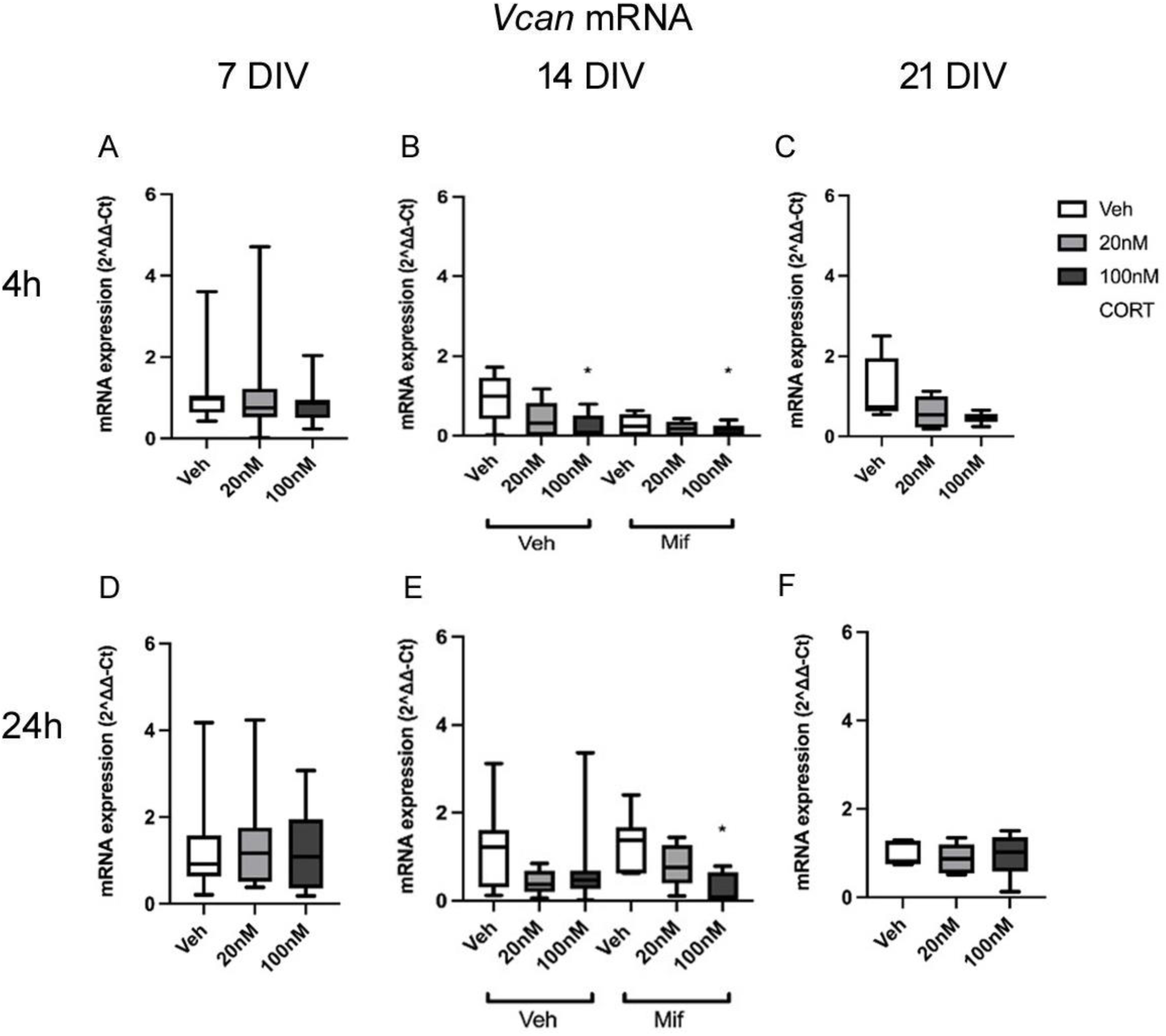
A-F: mRNA expression of *Vcan* after 4h and 24h exposure to low dose (20 nM) and high dose (100 nM) CORT at 7, 14 and 21 DIV, at 7 and 21 DIV with CORT alone, and at 14 DIV also with mifepristone (20 nM) treatment. Sample numbers: A,D - n=14-16/group; B-F - n=7-8/group. *p<0.05 vs corresponding vehicle group, post-hoc Tukey’s test. Boxes show median and interquartile range, with whiskers from minimum to maximum.

However, clear changes were observed at 14 DIV. There was a significant effect of CORT treatment at both 4h (F (2, 46)=3.32, p=0.046) and 24h (F(2, 46)=3.92, p=0.028) (Fig.1 B,E). The post-hoc testing revealed that the expression of *Vcan* decreased after high dose CORT treatment at both 4h and 24h relative to vehicle treatment (4h: veh vs high dose CORT p=0.048, 24h: veh vs high dose CORT p=0.021). The overall effect of mifepristone, independent of whether CORT was absent or present, was not significant at 4h (F(2, 46)=2.37, p=0.131) or 24h (F(2, 46)=0.01, p=0.943). However, the interaction of CORT and mifepristone was significant at 4h (F(2, 46)=4.63, p=0.016) (Fig. 1B), but not 24h (F(2, 46)=0.43, p=0.653) (Fig.1E), suggesting that mifepristone treatment exacerbated the corticosterone-driven suppression of *Vcan* expression at 4h (e.g. veh/mifepristone vs veh/veh - p=0.027 Fisher post-hoc test). Hence, glucocorticoids seem to exert a rapid non-GR-mediated suppression of *Vcan* mRNA levels, and a basal glucocorticoid receptor-mediated enhancement by GCs in the medium, which is rapidly blocked by mifepristone. In short, a rapid suppressive non-GR action and a rapid enhancing GR action on *Vcan* expression.

No significant effect of CORT on *Vcan* expression was observed at 21 DIV (Supplementary figure S3).

There was a tendency for *Acan* mRNA expression at 7 DIV to decrease after high dose CORT treatment at 4h (F (2, 43)=3.06, p=0.058; high dose VS vehicle, P=0.046, post-hoc Tukey test), but not 24h (F(2, 47) =1.32, p=0.277) (Fig.2 A,C). No significant effects were detected at 14 (Fig.1 B,D) or 21 DIV (Supplementary figure S3) (14 div 4h: F(2, 19)=2.11, p=0.152, 24h: F(2, 21)=2.11, p=0.152; 21 div 4h: F (2, 18)=1. 06, p=0.370, 24h: F (2, 19) =3.49, p=0.085), although the same trend for suppression at 4 but not 24h was noted at 14 DIV.

**Figure 2:**
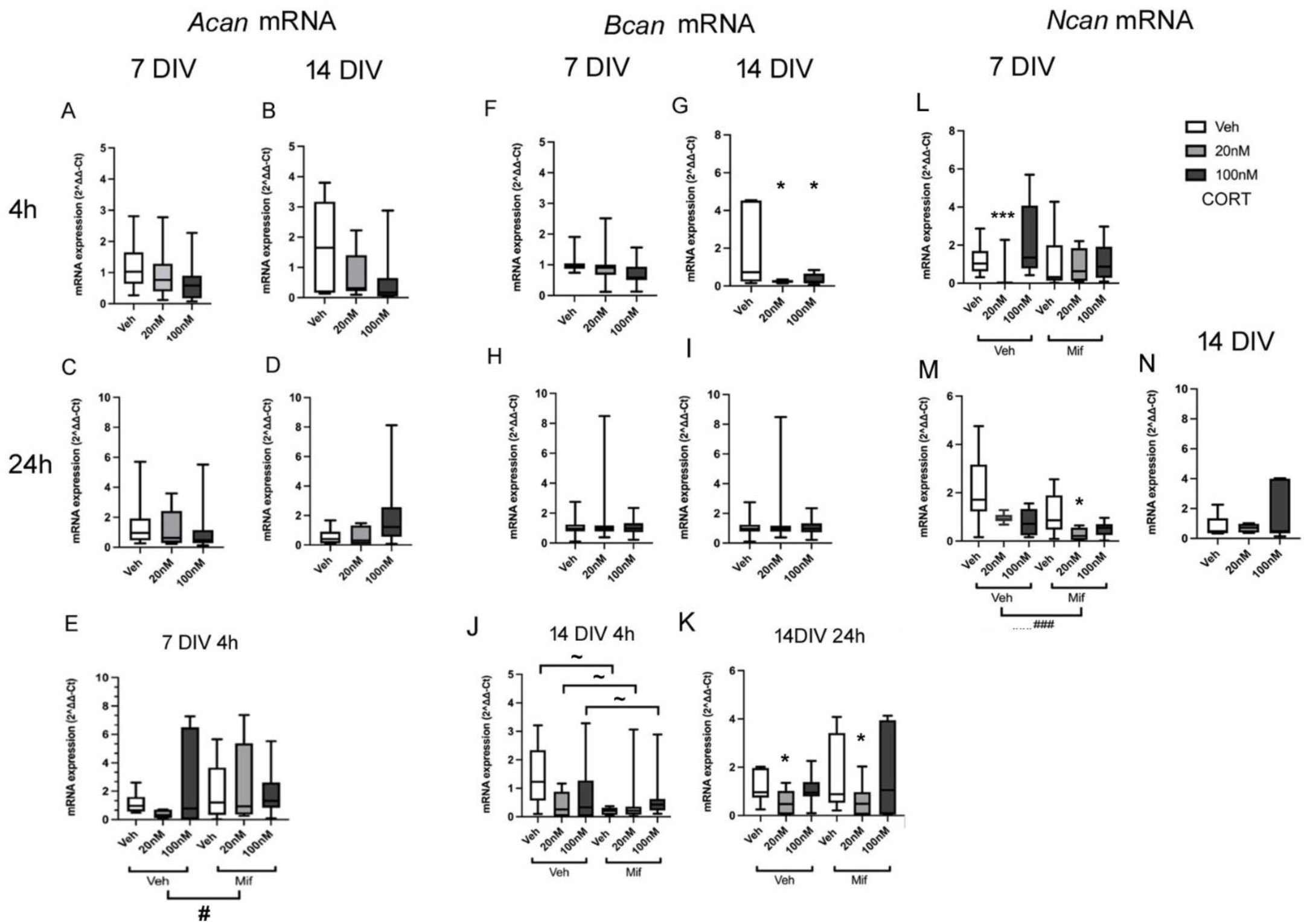
mRNA expression of *Acan* (A-E), *Bcan* (F-K), or *Ncan* (L-N) after exposure to vehicle (Veh), low dose (20nM) or high dose (100nM) CORT at 7 (A,C,E,F,H,L,M.) or 14 (B,D,G,I,J,K,N), with drug exposure for 4h (A,B,E,F,G,J,L) or 24h (C,D,H,I,K,M,N). 7 DIV experiments with CORT only, n=14-16/group; experiments also with mifepristone (20nM), n=7-8/group; 14 DIV experiments with CORT only, n=6-8/group; experiments also with mifepristone (20nM), n=7-8/group; 21 DIV n=6-8/group. # p<0.05, ### p<0.001 overall effect mifepristone vs vehicle groups (ANOVA); * p<0.05, *** p<0.001 vs corresponding vehicle group, ∼ p<0.05 for comparison shown, (post-hoc Tukey test). Boxes show median and interquartile range, with whiskers from minimum to maximum.

**Figure 3:**
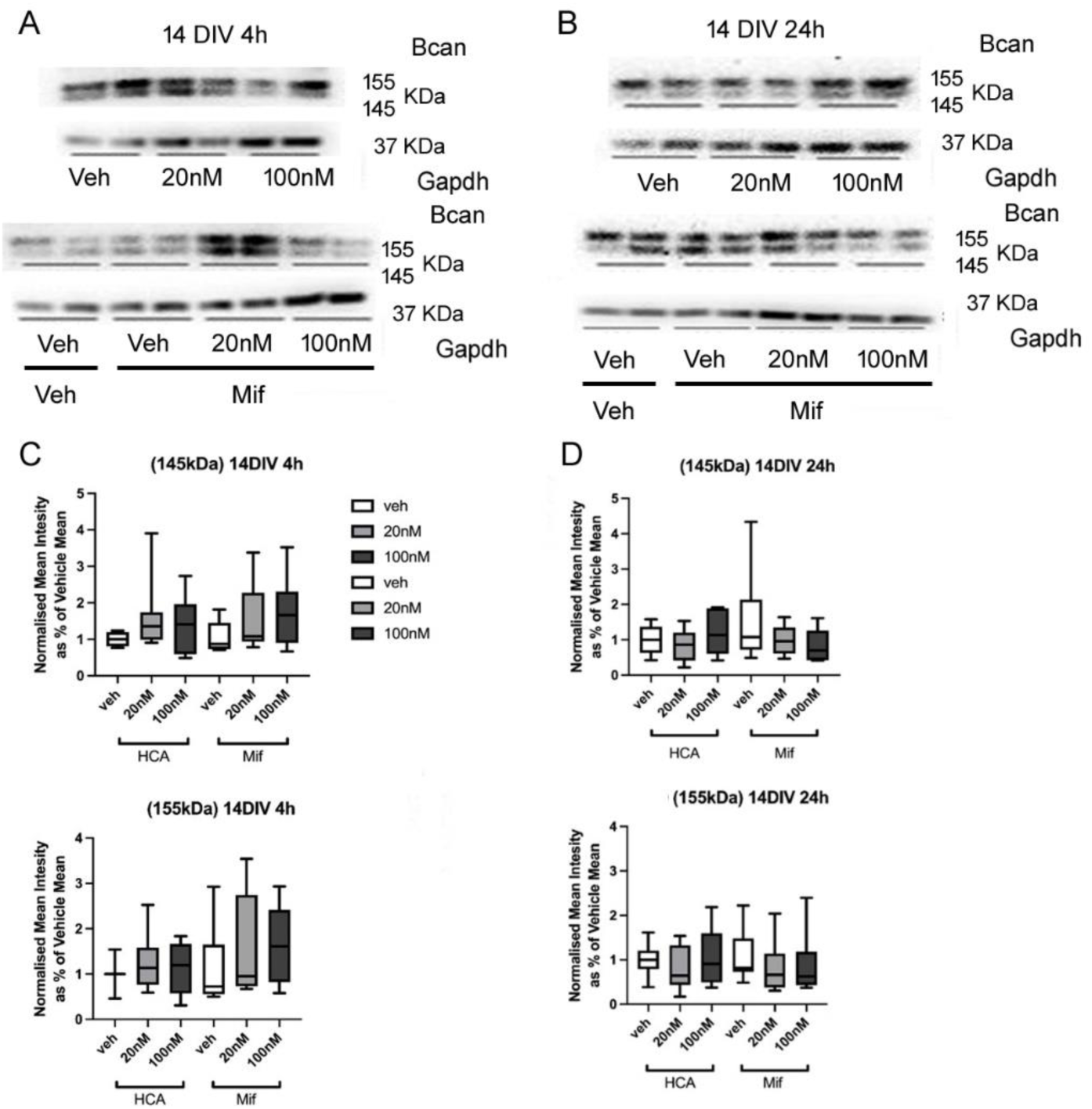
Bcan protein expression after exposure to CORT (20 or 100nM) and/or mifepristone (20nM). Representative results from western blotting are shown after 4h or 24h (**B**) exposure at 14 DIV. (**C,D**) corresponding box and whisker plots showing normalised mean intensity of the band signals after CORT and mifepristone exposure at 14 DIV with 4h (**C**) or 24h (**D**) exposure, for immunoreactive bands at 145 and 155 KDa. (n=46 in total, Veh: veh=7, 20nM=8, 100nM=8; Mifepristone: veh=7, 20nM=8, 100nM =8). Boxes show median and interquartile range, with whiskers from minimum to maximum.

When the experiment was repeated to test any influence of mifepristone, no significant effect of CORT was shown with 4h exposure (F (2, 44)=0.25, p=0.779) (Fig.2 E), although there was a hint of a suppression as before, (p=0.085 veh without mifepristone vs 20nM CORT without mifepristone, post-hoc Tukey test), but mifepristone overall significantly increased expression of *Acan* (F (2, 44)=4.18, p=0.048). The interaction of mifepristone and CORT treatment was not significant (F (2, 44)=1.78, p=0.183). The results suggested that there was some basal suppression of *Acan* expression by GCs, which limited the ability to detect further suppression with CORT application, but which was revealed by mifepristone exposure.

*Bcan* mRNA expression decreased significantly after 4h exposure to low dose and high dose CORT treatment at 14DIV (F (2, 19) =3.62, p=0.048; veh vs low dose: p=0.025, veh vs high dose p=0.035, Fisher post-hoc tests), despite considerable variability in the control group (Fig.2 G). No significant changes in gene expression were detected at 7DIV (4h: F (2, 43) =2.80, p=0.072; 24h: F (2, 47) =0.85, p=0.436) (Fig.2 F), 14DIV (24h: F (2, 21) =0.53, p=0.596)(Fig 2I) or 21 DIV (4h: F (2, 18) =1.06, p=0.704) (Supplementary Fig. S3). When the experiment was repeated in the absence or presence of mifepristone, *Bcan* mRNA levels were again found to be decreased by CORT in the absence of mifepristone (veh vs low dose CORT p=0.039, veh vs high dose CORT p=0.020, Fisher post-hoc tests). There was an overall significant effect of CORT treatment on *Bcan* mRNA expression at 14DIV after 24h (F (2, 46) =6.85, p=0.003), but not 4h (F (2, 46) =1.19, p=0.316), suggesting that the mRNA expression of *Bcan* decreased significantly after 24h low dose CORT treatment (veh vs low dose CORT p=0.002 Tukey post-hoc tests) (Fig. 2J,K). The results did not show a significant overall effect of mifepristone on *Bcan* expression, at either 4h (F (2, 46)=0.29, p=0.592) or 24h (F(2, 46)=0.97, p=0.329). These results indicated that at 14DIV, the ability of 20nM CORT to reduce *Bcan* mRNA levels was very clearly unaffected by the presence of mifepristone (Fig. 2J,K). There was a significant interaction of CORT and mifepristone treatment at 4h (F (2,46) =3.63, p=0.036), but not 24h (F(2, 46)=0.51, p=0.603). Mifepristone caused a suppression of *Bcan* mRNA levels (veh with mifepristone vs veh without mifepristone p=0.018, Fisher post-hoc tests). At 21DIV, the expression of *Bcan* was not affected by CORT at 4h (F(2, 44)=0.59, p=0.560) or 24h (F(2, 44)=2.78, p=0.075). Moreover, the effect of mifepristone was not significant either, at 4h (F(1,44)=0.01, p=0.936) or 24h (F(1, 44)=1.58, p=0.217). Similarly, the interaction of mifepristone and CORT treatment was not significant with 4h (F(1, 44)=1.24, P=3.000) and 24h (F(1, 44)=0.78, p=0.466) exposure (Supplementary figure 3).

Hence, just as with the regulation of *Vcan* expression, GCs seem to exert a rapid non-GR-mediated suppression of *Bcan* mRNA levels by CORT. These complex effects on *Bcan* mRNA suggest a rapid non-GR-mediated suppression by CORT, and a basal GR-mediated enhancement by GCs in the medium, which is rapidly blocked by mifepristone at 14DIV.

CORT exposure also showed an ability to suppress *Ncan* expression. At 7 DIV, a significant effect of CORT on *Ncan* was detected after 4h (Fig. 2L) (F(2,45)=7.66, p=0.002) and 24h (F(2, 45)=4.96, p=0.012), (veh vs low dose CORT at 4h p=0.008) and decreased with both low and high dose treatment at 24h (veh vs low dose CORT p=0.017, veh vs high dose CORT p=0.036, Tukey post-hoc tests). The same reduction was also detected in the presence of mifepristone, with lower expression after 24h low dose CORT (veh vs low dose CORT, p=0.023, Tukey post-hoc tests), but no significant changes with 4h CORT exposure in the presence of mifepristone (veh vs low dose CORT, p=0.869, veh vs high dose CORT, p=0.683, Tukey post-hoc tests). The interaction of mifepristone and CORT treatment was significant after 4h (F(2, 45)=9.79, p<0.001), and 24h (F(2, 45)=2.20, p=0.124), in that expression of *Ncan* with 4h low dose CORT exposure was increased in the presence of mifepristone compared to in its absence (p=0.024, Tukey post-hoc tests), but decreased for the same comparison after 24h (p=0.006). Additionally, there was an overall effect of mifepristone to decrease *Ncan* mRNA expression over 24h (F(2, 45)=12.47, p=0.001), but not 4h (F(2,45)=1,06, p=0.310). Overall this suggests a suppressive effect of slightly increasing GC levels at 4h through GRs (mifepristone sensitive), and also not via GRs over a longer time scale (mifepristone insensitive), combined with the basal levels of GCs in the medium tending to enhance *Ncan* expression (probably genomic in mechanism, since slowly relieved over 24h by mifepristone, resulting in a further decrease in levels).  No changes in Ncan expression were detected at 21 DIV (Fig. 2 O, Supplementary figure 3).

Although several *Bcan* mRNA alterations were observed at 14DIV, the protein expression of *Bcan* (155kDa) (Fig.2 K L) (4h: F (2, 26)=1.5, p=0.227) remained unchanged after 4h CORT exposure. With mifepristone cotreatment, no significant changes were observed (Fig.2 L M) (155kDa/145kDa) protein levels with both 4h and 24h CORT exposure (155kDa: 4h F(2, 47)=0.97, p=0.389, 24h: F(2, 47)=0.77, p=0.469; 145kDa:4h F(2, 47)=2.64, p=0.083, 24h: F(2, 47)=1.10 p=0.342). Moreover, there were no overall effects of mifepristone (155kDa: 4h F(1, 47)=1.54, p=0.221, 24h: F(1, 47)=0.00, p=0.963; 145kDa:4h F(1, 47)=0.21, p=0.652, 24h: F(1, 47)=0.23, p=0.631) and no significant interactions of CORT and mifepristone (155kDa: 4h F(2, 47)=0.27, p=0.765, 24h: F(2, 47)=0.28, p=0.759; 145kDa:4h F(2, 47)=0.38, p=0.686, 24h: F(2, 47)=1.73, p=0.190) detected after 4h and 24h exposure.

For *phosphacan* (*Ptprz1*), mRNA expression was unchanged after 4h or 24h CORT exposure, with low or high doses, at 7 DIV, 14 DIV and 21 DIV (Supplementary figures S3, S4).

*Has1* gene expression was not significantly affected at 7 DIV (4h: F (2, 21) =2.02, P=0.161; 24h: F (2, 23) =2.71, p=0.09) (Fig.4 A) or 14 DIV (4h: F (2, 8) =0.07, p=0.933, 24h: F (2, 12) =1.75, p=0.223) (Fig.4 C and data not shown), although there was a trend towards a decrease in *Has1* expression at 7 DIV after 4h with 100nM CORT (p=0.068, post-hoc Tukey test). When repeated without or with mifepristone, there was again a tendency towards decreased *Has1* mRNA expression with 4h CORT exposure to the lower dose (Fig.4 A, B), but there was also a great deal of variability with the higher dose. Overall, including mifepristone groups, there was no significant CORT effect (F (2, 44) =0.204 p=0.959) (Fig.4 B). There was however an overall effect of mifepristone (F (2, 44) =6.75, p=0.014), indicating mifepristone significantly increased expression of *Has1*, consistent with medium GCs acting to suppress mRNA levels. The mifepristone x CORT treatment interaction was not significant (F (2, 44) =1.78, p=0.183). After long-term (24h) exposure to CORT and mifepristone at 7 DIV, the mRNA levels of *Has1* still remained unchanged (F (2, 44) =1.34, P=0.273), and with no overall effect of mifepristone (F (1, 44) =1.29, p=0.263) and no interactions of CORT and mifepristone (F (2, 44) =0.26, p=0.775) were found (data not shown). At 14DIV, no significant effect of CORT was observed after 24h CORT treatment (F(2, 46)=0.88, p=0.422), but there was now a really clear overall effect of mifepristone (F(2, 46)=12.48, p=0.001) (Fig 4C). The interaction of mifepristone and CORT treatment was not significant (F(2, 46)=0.35, p=0.708). These results suggested that mifepristone significantly blocked a basal suppression of *Has1* expression by glucocorticoids in the culture medium, probably rendering it problematic to reveal a clear additional effect of CORT addition.  The results suggested a rapidly relievable effect of glucocorticoids in the culture medium to suppress *Has1* mRNA levels at 7 and 14 DIV, an effect attenuated by mifepristone exposure, and slightly enhanced by exogenous CORT exposure. 

**Figure 4:**
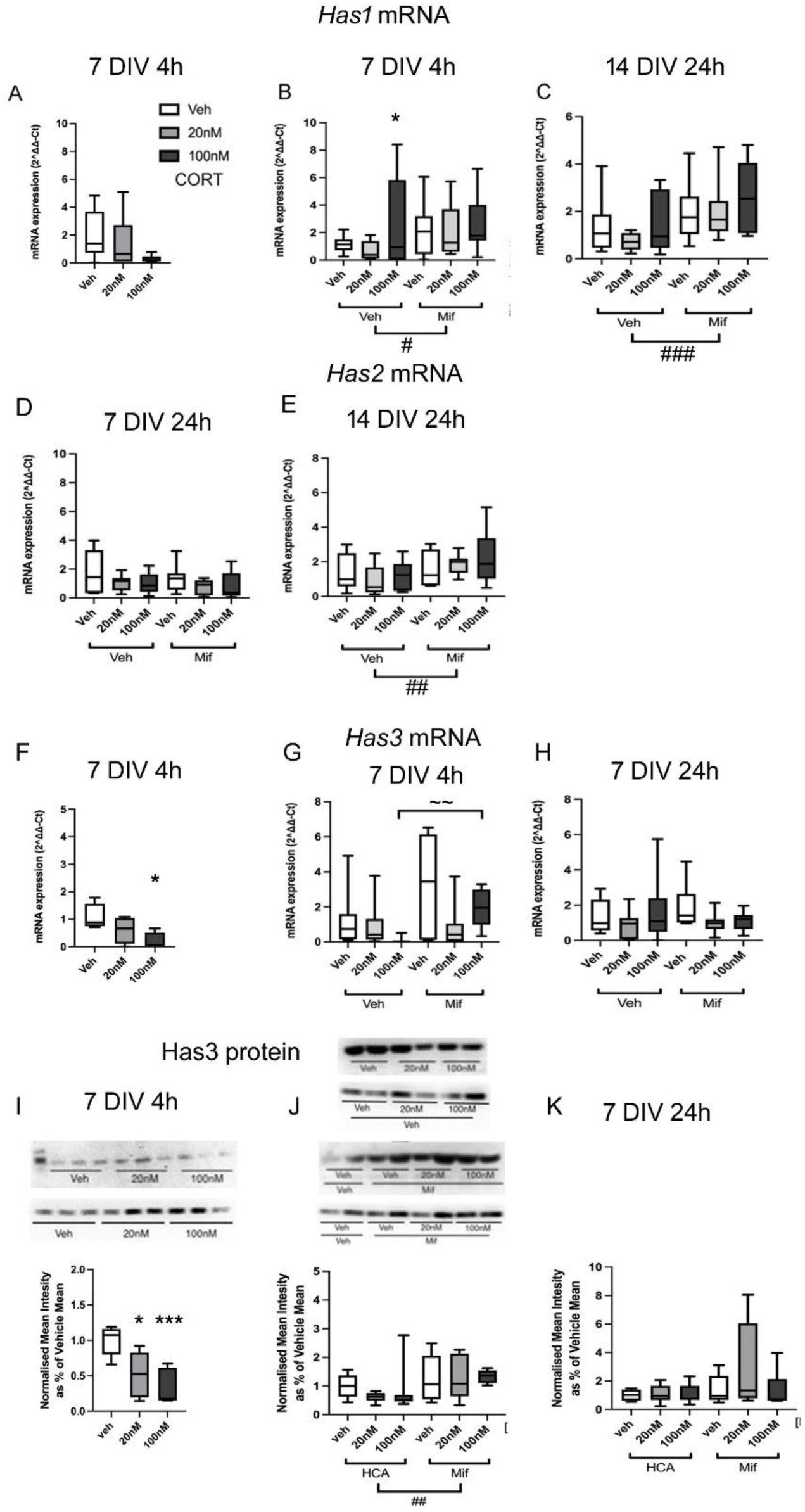
Effects of CORT on Has gene expression **A-C**: *Has1* mRNA expression; **D,E**: *Has2* mRNA expression, and **F-H**: *Has3* mRNA expression after 4h or 24h exposure to low dose (20nM) or (100nM) high dose CORT at 7 or 14 DIV, in the absence or presence of mifepristone (20nM). (7 DIV, n=6-8/group; 14 DIV, n=7-8/group; **I,J**: Western blot analysis of Has3 protein levels at 7 and 14 DIV after 4h or 24h exposure to 20nM or 100nM CORT alone (**I**), or without or with mifepristone (20nM) treatment (**J**). n: veh=8, 20nM =8, 100nM =8; Mifepristone: veh=8, 20nM=8, 100nM =8. Example blots are shown, with corresponding quantification below. **K**: Corresponding western blot analysis of Has3 protein levels exposed to the same doses of COR for 24h. n: veh=12, 20nM CORT=12, 100nM CORT=10. *p<0.05, ***p<0.001 vs corresponding vehicle group, post-hoc Tukey’s test; ∼∼ p<0.01 for comparison shown, post-hoc Fisher’s test; # p<0.05, ## p<0.01, ### p<0.001, overall effect of mifepristone vs corresponding (mifepristone) vehicle group, ANOVA. Boxes show median and interquartile range, with whiskers from minimum to maximum.

For *Has2* mRNA expression, no overall effect of mifepristone was detected at 7 DIV at 4h (F (2, 44) =2.50, P=0.123), with no significant interaction of mifepristone and CORT treatment (F (2, 44) =0.80, p=0.457) (data not shown). Equally, no alterations of *Has2* mRNA levels were observed after longer-term exposure to CORT (F (2, 44) =1.98, p=0.151) at 7 DIV, and no overall effect of mifepristone was detected either (F (1, 44) =1.85, p=0.181) (Fig.4 D), with no interactions between CORT and mifepristone (F (1, 44) =0.38, p=0.688. At 14 DIV, the changes in *Has2* expression were again not significant 24h after CORT treatment (F (2, 43) =0.36, p=0.701), but the expression of *Has2* was significantly affected by mifepristone (F (2, 43) =7.20, p=0.011) (Fig.4 E), suggesting increased *Has2* expression after blocking GRs. Interactions of mifepristone and CORT treatment were not significant (F (2, 43) =1.46, p=0.246). At 21 DIV, the effect of CORT was not significant at 24h (F (2, 46) =0.25, p=0.783) (Supplementary figure S3). Mifepristone did not have an overall effect on *Has2* expression at 24h either (F (1, 47) =0.11, p=0.738). And there was no significant interaction of mifepristone and CORT treatment detected with 24h (F (2, 46) =1.10, p=0.303) exposure. 

*Has3* mRNA expression decreased significantly at 7 DIV after 4h 100nM CORT treatment (F (2,9) =5.40, p=0.038, veh vs high dose CORT, p=0.039, Tukey post-hoc test) (Fig.4 F). With mifepristone, at 7 DIV, while there was no significant effect of CORT treatment overall at either 4h (F(2,45)=0.82, p=0.452) or 24h (F(2,45)=1.97, p=0.753) (Fig.4 G,H), and the overall effect of mifepristone was not significant either, at 4h (F(2,45)=2.04, P=0.163) or 24h (F(2,45)=2.45, P=0.126) (Fig.4 G,H), there was a significant interaction between CORT and mifepristone treatment after 4h (F(2, 45)=3.65, p=0.039), but not 24h (F(2, 45)=0.05, p=0.955), suggesting a tendency for mifepristone to attenuate the suppression of *Has3* mRNA levels by exposure to the high dose of CORT (high dose CORT with mifepristone vs high dose CORT without mifepristone p=0.009, Fisher post-hoc test). At 21 DIV, no significant effects of CORT on the expression of *Has3* mRNA levels were found at 4h (F(2,47)=1.79, p=0.179) or 24h (F(2, 47)=0.24, p=0.787) (Supplementary figure S3). There was no effect of mifepristone at 4h (F(1, 47)=0.30, p=0.43) or 24h (F(1,47)=2.70, p=0.108). Additionally, no significant interaction of mifepristone and CORT treatment was detected with 24h (F(2,47)=0.30, P=0.743) exposure (Supplementary figure S3). The results imply that, at 7 DIV, a suppressive effect of CORT on *Has3* mRNA levels is rapidly relieved by antagonism of GRs. 

We checked for corresponding protein alterations after CORT exposure, Has3 protein levels decreased significantly after 4h exposure with both low dose and high dose CORT (F(2,17)=11.14, p=0.014 veh vs low dose CORT, p=0.001 veh vs high dose CORT, Tukey post-hoc test) (Fig 4I). When re tested with mifepristone, the same trend was observed for CORT in the absence of mifepristone at 4h but not 24h exposure, but no overall CORT effect was found at either 4h (F (2, 47) =0.41, p=0.665) or 24h (F (2, 47) =1.21, p=0.309) of CORT and mifepristone exposure (Fig.4 J,K). However, mifepristone significantly increased *Has3* protein levels after 4h (F (1, 47) =8.29, p=0.006) (Fig.4 J), but not quite significantly after 24h exposure (F (1, 47) =3.38, p=0.073) (Fig.4 K). No significant interactions of CORT and mifepristone were observed with either 4h or 24h exposure (4h F (2, 47) =0.51, p=0.603, 24h: F (2, 47) =1.12, p=0.336). The evidence therefore points to a rapid suppression of neuronal *Has3* expression at both mRNA and protein levels by GCs via GRs.

Expression of *Hapln4* mRNA was unaffected by CORT after either dose or treatment time, except for a clear suppression of mRNA levels at 4h (but not 24h) after treatment at 14 DIV that was not attenuated by mifepristone (Supplementary figure S5). However, Hapln4 protein levels were unchanged by CORT exposure at 14 DIV (Supplementary figure S5).

The mRNA levels for *TnR* decreased significantly at 7 DIV after 4h low dose CORT treatment (F (2, 45)=3.28, p=0.049, veh vs low dose CORT p=0.038, Tukey post-hoc tests), but while the same trend was observed at 24h, the effect was not significant (F(2,45)=0.49, p=0.619). This decrease was maintained in the presence of mifepristone after 4h CORT exposure, but the mRNA level decreased with the presence of mifepristone with high dose CORT. There was an overall effect of mifepristone on *TnR* expression which was significant after 24h (F (2, 45)=9.17, P=0.004), but not 4h (F(2, 45)=1.69, P=0.201), (Fig.5 D). The level of mRNA expression with 24h mifepristone and CORT exposure was lower than that in the absence mifepristone.

**Figure 5:**
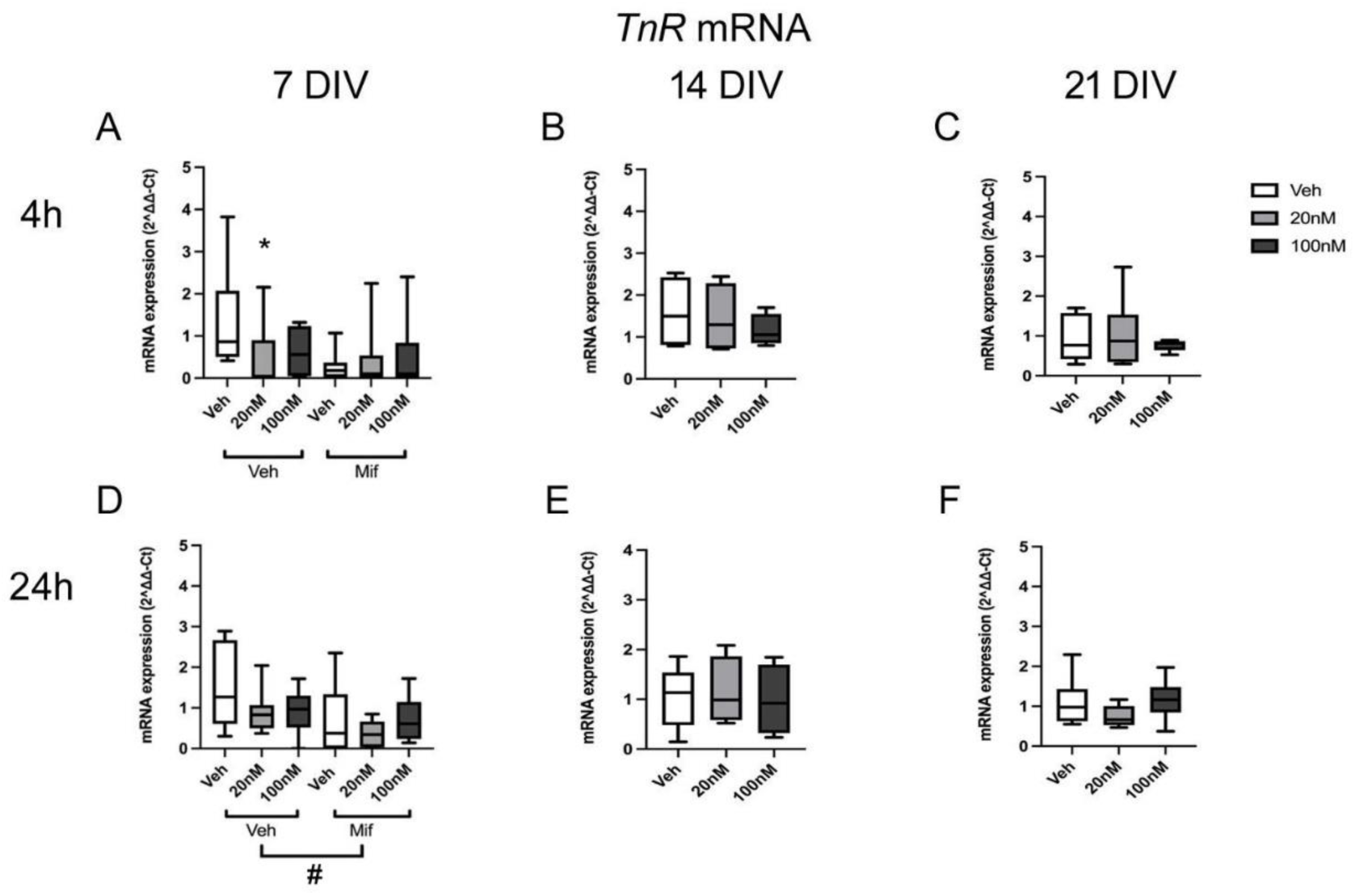
**A-F**: mRNA expression of *TnR* after 4h or 24h exposure to low dose (20nM) or high dose (100nM) CORT at 7, 14 and 21 DIV (B-F), and in the absence or presence of mifepristone (20nM) (A,D). (7 DIV: n=7-8/group; 14 DIV: n=9-11/group; 21 DIV: n=12-14/group). # p<0.05, overall effect of mifepristone vs corresponding (mifepristone) vehicle group, ANOVA. Boxes show median and interquartile range, with whiskers from minimum to maximum.

In addition, there was a significant interaction of CORT treatment and mifepristone treatment after 24h (F(2, 45)=5.42, P=0.008) rather than 4h (F(2, 45)=2.51, P=0.095), suggesting a reduction of mRNA levels of veh treatment group after long-term mifepristone exposure (veh without mifepristone vs veh with mifepristone p<0.001, low dose CORT with mifepristone vs low dose CORT without mifepristone p=0.045, Fisher post-hoc tests). There were no significant changes in *TnR* expression after CORT exposure at 14 DIV (4h: F (2, 12) =1.32, P=0.309; 24h: F (2, 11) =0.33, P=0.729) (Fig.5 B,E) or 21 DIV (4h: F (2, 16) =0.01, P=0.990; 24h: F (2, 20) =1.65, P=0.219) after either 4h and 24h treatment (Fig.5 C,F).

In contrast to the decrease in mRNA expression, protein levels of *TnR* remained unchanged with 4h or 24h CORT exposure at 7 DIV (Supplementary figure S6).

### CORT alters GABAergic gene expression

As PNNs surround mainly Pvalb-expressing GABAergic interneurons (Gabungcal et al., 2013; Morishita et al., 2015), GABA related components, including glutamate decarboxylase (*Gad*) genes and *Pvalb w*ere also measured in the current study.

The expression of *Pvalb* decreased significantly after 24h high dose CORT treatment at 21 DIV (F(2, 20)=4.72, p=0.023, veh vs high dose CORT p=0.043, Tukey post-hoc tests), but not 4h (F(2, 18)=0.26, p=0.775) (Fig.6 C,F). However, there were no significant changes at 7 DIV (4h: F(2, 23)=1.25, p=0.308; 24h: F(2, 22)=0.50, p=0.613) (Fig.6 A,D) or 14 DIV (4h: F(2, 23)=0.25, p=0.785; 24h: F(2, 22)=1.31, p=0.292) (Fig.6 B,E), after either 4h or 24h CORT treatment.

**Figure 6:**
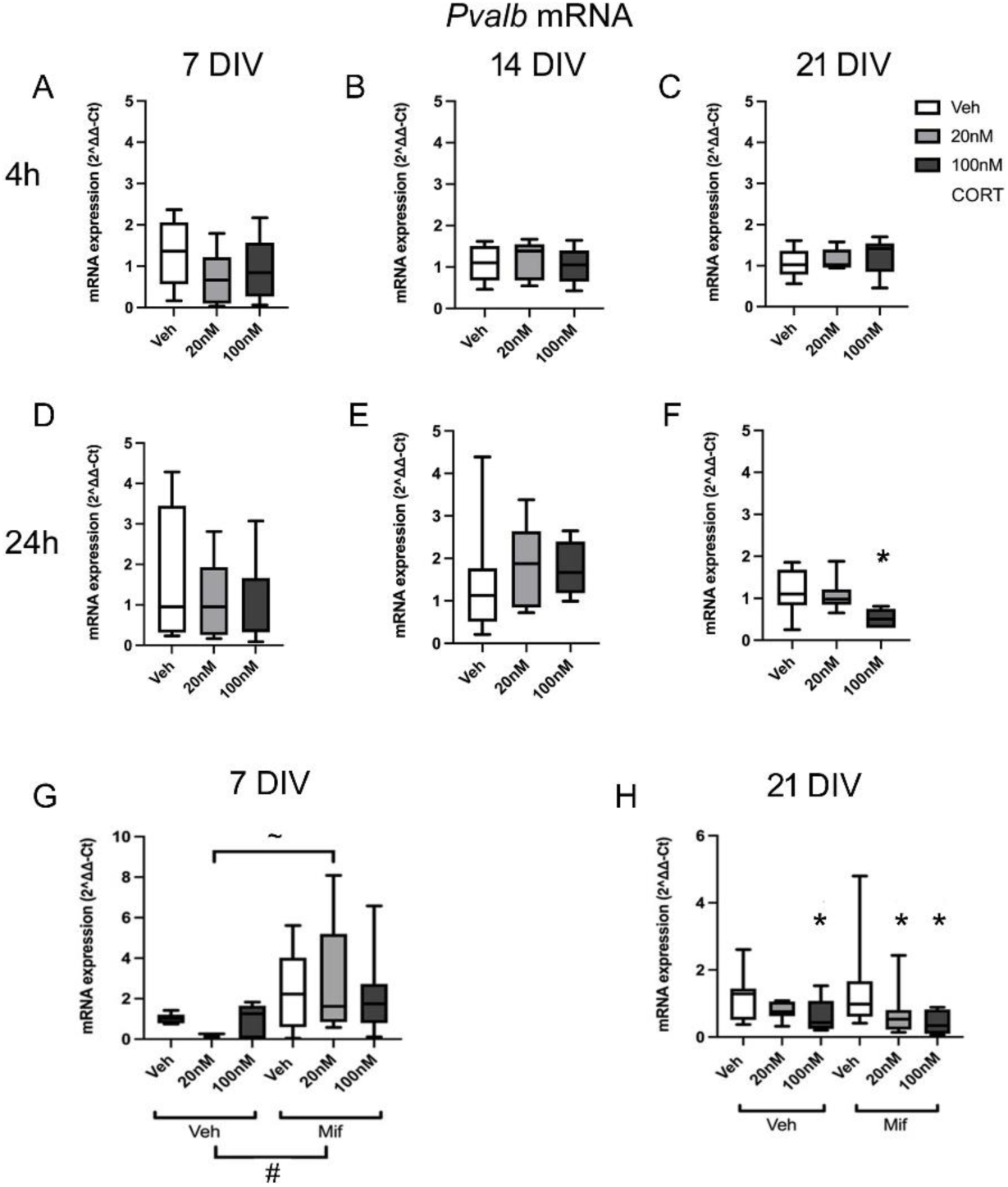
**A-F**: mRNA expression of *Pvalb* after 4h and 24h exposure to low dose (20nM) or high dose (100nM) CORT at 7, 14 and 21 DIV. (7 DIV: n=20 in total, veh=8, low dose=6, high dose=7; 14 DIV: n=28 in total, veh=9, low dose=11, high dose=9; 21 DIV: n=39 in total, veh=14, low dose=14, high dose=12). **G-H**: mRNA expression of *Pvalb* at 7 DIV (4h) and 21 DIV (24h), after CORT and mifepristone (20nM) treatment. (n=45 in total, Veh: veh=8, low dose samples=8, high dose samples=7; Mifepristone: veh=8, low dose samples=7, high dose samples=7) *p<0.05 vs corresponding vehicle group, ∼ p<0.05 for comparison shown, post-hoc Tukey’s test; # p<0.05 overall effect of mifepristone vs corresponding vehicle group, ANOVA. Boxes show median and interquartile range, with whiskers from minimum to maximum.

When CORT treatment was repeated, either in the absence or presence of mifepristone, the results again showed that at 7 DIV, there was no significant effect of CORT on the expression of *Pvalb* after 4h exposure (F(2, 43)=0.89, p=0.420) (Fig. 6 G). However, there was a significant overall effect of mifepristone (F(1, 43)=12.40, p=0.001) with an increase in expression of *Pvalb* in cultures exposed to mifepristone. For individual group comparisons, the *Pvalb* mRNA levels significantly increased with 4h exposure by 20nM CORT with mifepristone compared to 20nM CORT without mifepristone (p=0.022, Tukey post-hoc tests), with the same trend at other CORT doses. This implies a basal GR-mediated suppression of *Pvalb* mRNA at 7 DIV (despite the low basal levels of expression at this developmental stage). At 21 DIV, there was a once more a significant effect of high dose CORT after 24h exposure, maintained in the presence of mifepristone (F(2,47)=8.66, p=0.001; veh with mifepristone vs low dose CORT with mifepristone p=0.035, veh with mifepristone vs high dose CORT with mifepristone p<0.001, post-hoc Tukey test) (Fig.6 H). The overall effect of mifepristone was not significant (F(1,47)=1.88, p=0.178), and the interaction of mifepristone and CORT treatment was not detected with 24h exposure (F(2,47)=0.88, p=0.423), suggesting that the clear suppressive effect of CORT on *Pvalb* expression was not mediated by GRs.

*GAD1* expression is also robustly found to be decreased in PFC in schizophrenia (Gonzalez-Burgos et al., 2015; Hoftman et al., 2015). When tested either in the absence or presence of mifepristone at 7 DIV, the expression of *Gad1* was not significantly changed after 4h CORT exposure (F (2,40)=0.93, p=0.406) (Fig.7A). Equally, no changes were observed after 24h exposure (Supplementary figure S7). However, the overall effect of mifepristone after 4h exposure approached significance (F (1,40) =4.01, p=0.053), suggesting a possible suppression of *Gad1* expression by GCs in the culture medium that is relieved in the presence of mifepristone. There was no significant interaction between CORT and mifepristone exposure (F (1,40) =4.01, p=0.237). At 14 DIV, after 4h exposure, no significant overall change in *Gad1* mRNA levels was found for either CORT (F (2,47) =0.22, p=0.803) or mifepristone (F (1,47) =0.19, p=0.661) (Fig.7A). However, there was a significant interaction of CORT and mifepristone (F (2,47) =0.823, p=0.038), with reduction of mRNA levels of *Gad1* after high dose (100nM) CORT exposure relative to vehicle in the absence of mifepristone (p=0.045, Fisher post-hoc test) (Fig. 7 A). At 21 DIV, with 24h exposure, no effect of either CORT (F (2,46) =0.71, p=0.496) or mifepristone (F (1,46) =0.21, p=0.647) was observed, with no significant interaction between the two treatments (F (2,46) =0.23, p=0.798) (Fig 7A).

**Figure 7:**
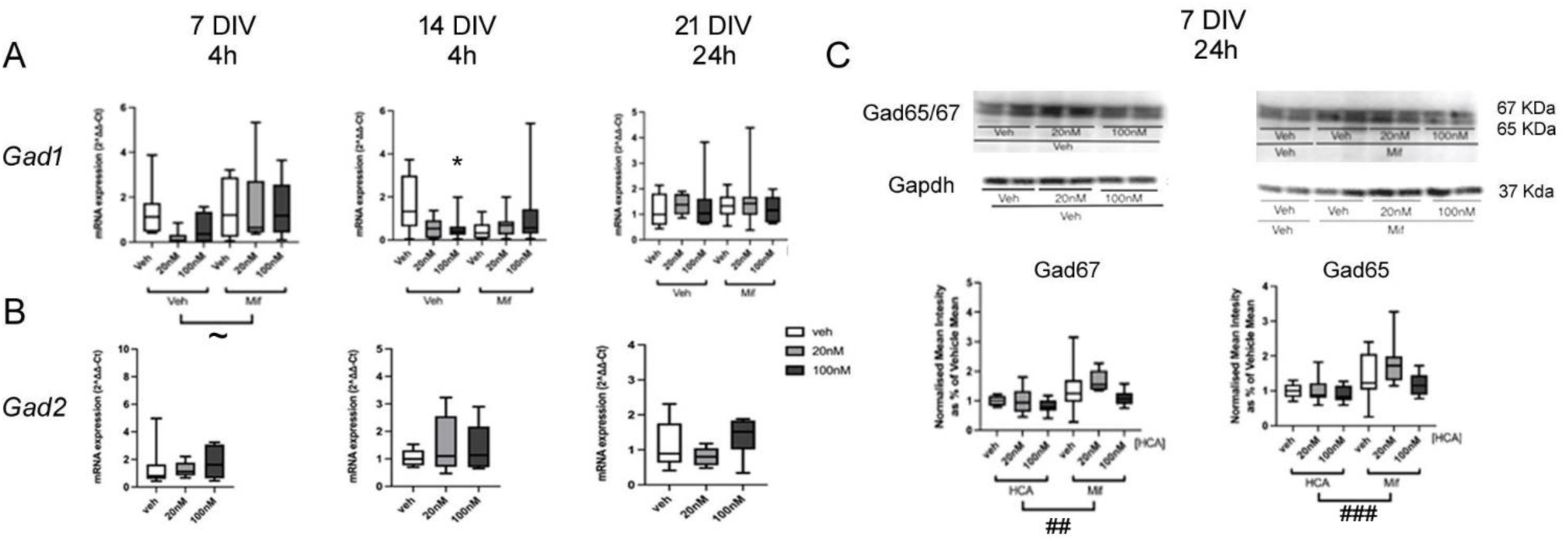
**A,B**: mRNA expression of *Gad1* (A) and *Gad2* (B) after 4h (left and centre graphs) and 24h (right graph) exposure to low dose (20nM) or high dose (100nM) CORT at 7, 14 and 21 DIV. n=7-8/group (GAD1), 6-8/group (Gad2 7 DIV), 9-11/group (Gad2 14 DIV) or 12-14/group (Gad2 21 DIV). **C**: Immunoblotting with anti-Gad65/67 and anti-Gapdh antisera at 7 DIV (24h) after CORT and mifepristone (20nM) treatment. (n=96 in total: veh: veh=16, low dose CORT=16, high dose CORT=16; Mifepristone: veh=16, low dose CORT=16, high dose CORT=16). * p<0.05 vs corresponding vehicle group (post-hoc Fisher’s test); ∼p=0.053, ## p<0.01, ### p<0.001, overall effect of mifepristone vs corresponding vehicle groups, ANOVA. Boxes show median and interquartile range, with whiskers from minimum to maximum

In contrast to the effects of CORT on *Gad1* expression, no changes were detected in *Gad2* (GAD65) mRNA expression following CORT exposure at 7, 14 or 21 DIV at 4 or 24h (F(2,23)=0.38, p=0.687 (4h); F(2,23)=0.33, p=0.723 (4h); F(2,20)=0.73, P=0.495 (24h) respectively) (Fig 7B, Supplementary figure S7).

Protein levels were also measured based on the mRNA alterations, at 7 and 14 DIV after 24h exposure. At 7 DIV, there were no significant changes of *Gad67/65* with 24h CORT exposure, although there was a slight trend towards decreased expression (*Gad67*: 24h F(2, 47)=3.15, p=0.053; *Gad65*: 24h: F(2, 47)=2.61, p=0.085) (Fig.7 C). However there was a significant overall effect of mifepristone on *Gad65/67* protein levels (*Gad67*: 24h F(1, 47)=12.72, p=0.001; *Gad65*: 24h: F(1, 47)=13.68, p=0.001), with increased expression at 24h. There were no significant interactions of CORT and mifepristone were detected inn *Gad67/65* (*Gad67*: F(2, 47)=0.72, p=0.493; *Gad65*: 24h: F(2, 47)=2.61, p=0.085) (Fig.7 C). At 14 DIV, neither CORT nor mifepristone had any significant effect on Gad1/Gad67 or Gad2/Gad65 protein levels after 24h exposure (Supplementary Figure S8). These results suggested that CORT had little overt effect on Gad1/Gad67 protein levels, however at 7 DIV, but not later in development, mifepristone exposure revealed a basal suppression of both Gad67 and Gad65 protein, and *Gad1* mRNA, by basal levels of GCs in the culture medium.

### Effect of Collagen 3 (GPR56/97 agonist) and aldosterone (MR agonist)

The previous results showed that glucocorticoids could regulate the expression of PNN components, and the changes of *Has3, Gad1*, and *Pv* at 7 DIV could be reversed by the selective GR antagonist, mifepristone. However, the expression of other PNN components which was significantly regulated by glucocorticoids was not reversed by mifepristone. In this case, the effect might be activated via other GC targets rather than via GRs. Therefore, GPR56/GPR97 (ADGRG1/ADGRG3) were considered as a potential alternative pathway for the GC effect, as GCs are agonists at these receptors (GPR97/ADGRG3) (Y.-Q. Ping et al., 2021), collagen 3, an agonist of GPR97 and GPR56 (Luo et al., 2011; Olaniru et al., 2018; Zhu, Luo, et al., 2019) (considering the similarity between GPR97 and GPR56 at the binding site, they are likely to share the same ligands)(Vizurraga et al., 2020) was tested on PNN component mRNAs which were affected by GCs but were not sensitive to mifepristone reversal (7 DIV: *Ncan*, *TnR;* 14 DIV: *Bcan*, *Vcan*, *Hapln4*, *Gad1*).

The mRNA expression of *TnR* was significantly upregulated by collagen 3 treatment for 4h at 7DIV (P=0.020, F (1, 23) =6.33) (Fig 8 A). *Ncan* mRNA expression was downregulated by collagen 3 treatment for 4h (P=0.006, F (1, 23) =9.29) (Fig. 8 A) at 7DIV. The results indicated that collagen 3 could increase *TnR* mRNA expression and suppress *Ncan* mRNA expression rapidly. The former effect is opposite to the effect of GCs, but the effect on *Ncan* expression is similar, raising the possibility that both collagen 3 and GCs could be acting via the same (non-genomic) GPCR-mediated mechanism in this case.

**Figure 8:**
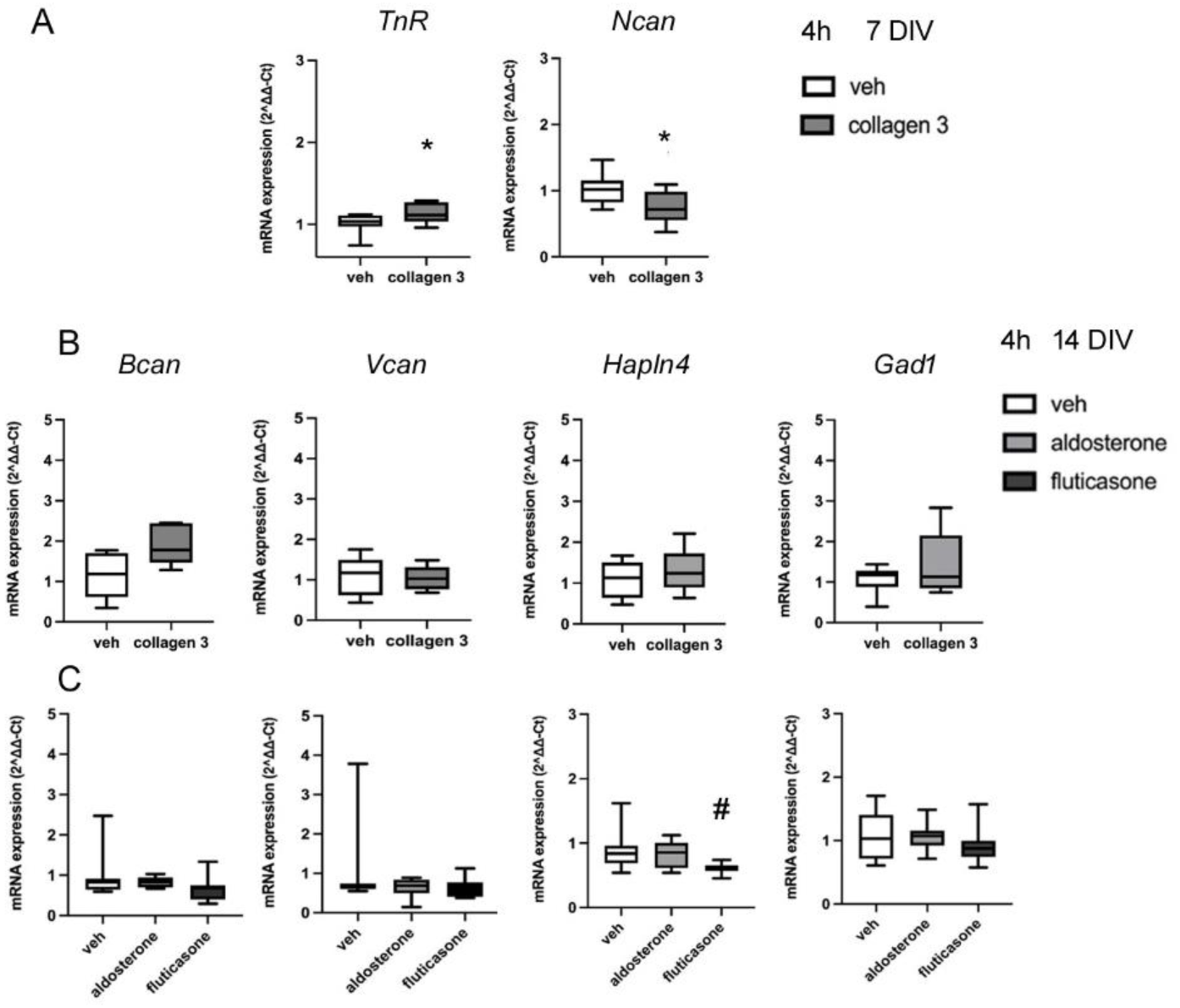
**A**. mRNA expression of *TnR* and *Ncan* at 7DIV after 4h collagen 3 (75nM) treatment; (n=23 in total, veh=12, collagen 3 treated samples=11). **B**. mRNA expression of *Bcan, Vcan, Hapln4* and *Gad1* at 14DIV after 4h collagen 3 (75nM) treatment. C. mRNA expression of *Bcan, Vcan, Hapln4* and *Gad1* at 14DIV after 4h treatment with aldosterone (100nM) or fluticasone (50nM). (n=22 in total, Veh=6, aldosterone treated samples=8, fluticasone treated samples=8).* p<0.05 vs vehicle group, ANOVA; # p=0.024 vs vehicle group, Mann Whitney U test. Boxes show median and interquartile range, with whiskers from minimum to maximum.

In addition, the mRNA levels of *Bcan* (P=0.071, F(1, 10)=4.20), *Vcan* (P=0.985, F(1, 10)=0.00), *Hapln4* (P=0.528, F(1, 10)=0.43) and *Gad1* (P=0.434, F(1, 10)=0.67) were not affected by collagen 3 at 14 DIV, suggesting that mRNA levels of *Bcan*, *Vcan, Hapln4* and *Gad1* were not affected through an action of glucocorticoids binding to GPR56/97 (Fig. 8 B).

In order to explore the mechanisms involved in these non-GR actions further, we compared the effects of the selective MR agonist aldosterone with the selective (no MR actions) GR agonist fluticasone. The mRNA expression of *Bcan* (P=0.938; F(2,22)=1.77), *Vcan* (P=0.440; F(2,22)=1.02), *Hapln4* (P=0.887; F(2,22)=3.30) and *Gad1* (P=0.0.981; F(2,22)=0.68) remained unchanged after exposure to aldosterone at 14DIV for 4h (Fig 8 C). In terms of the effect of fluticasone, there was a decreasing tendency of *Hapln4* mRNA expression (P=0.060; F(2,22)=1.77; Veh vs fluticasone, p=0.024, Mann Whitney U test) (Fig 8 C), suggesting an inhibition effect of fluticasone mediated by GRs. However, there was no significant change for *Bcan* (p=0.201; F(2,22)=1.77), *Vcan* (p=0.460; F(2,22)=1.02) or *Gad1* (p=0.636; F(2,22)=0.68) expression (Fig 8 C). These results indicated that MRs were almost certainly not the mediator of the effects of CORT on the expression of these genes. Further, the clear lack of effect of fluticasone in the cases of *Vcan* and *Gad1* provided additional evidence that GRs are also not involved in these effects.

### No evidence for GR-mediated mRNA decay involvement

The GRs in the cytoplasm reportedly can bind directly to RNA, and GCs can activate the RNA-bound GR, leading to mRNA degradation, defined as the GC-mediated mRNA decay (GMD) pathway (Cho et al., 2015). However, no alteration in the rate of *Bcan* or *Hapln4* mRNA degradation was detected after exposure to 20nM CORT (Supplementary figure S9).

### Glucocorticoids suppress proteasome activity

To test whether the glucocorticoids affect proteasome activities, neuronal cultures at 14DIV, treated with hydrocortisone for 4h, were measured with fluorogenic proteasome assay with 3 different proteasome substrates related to the 3 proteasome activities, including chymotrypsin-like activity, caspase-like activity, and trypsin-like activity. All activity was completely inhibited by 10μM MG132 (data not shown).

The results of the proteasome assays showed that low and high doses of CORT tended to inhibit the proteasome activities. The chymotrypsin-like activity could be inhibited by both low dose and high dose CORT compared to the vehicle group (Fig. 9), however, only the inhibition by high dose CORT was statistically significant (F (2, 11) =7.18, p=0.011 veh vs high dose CORT; p=0.144 veh vs low dose CORT, post-hoc Tukey test). However, no apparent effect of low or high dose CORT on caspase-like activity was observed (F (2, 11) =0.32, p=0.910 veh vs low dose CORT; p=0.923 veh vs high dose CORT, post-hoc Tukey test) (Fig 9). Moreover, only very low activity was observed using the substrate which detected the trypsin-like activity (data not shown). Although the inhibition tendency was shown with high dose CORT exposure, the inhibition was not significant (F (2, 11) =2.65, p=0.947 veh vs low dose CORT; p=0.219 veh vs high dose CORT, post-hoc Tukey test).

**Figure 9:**
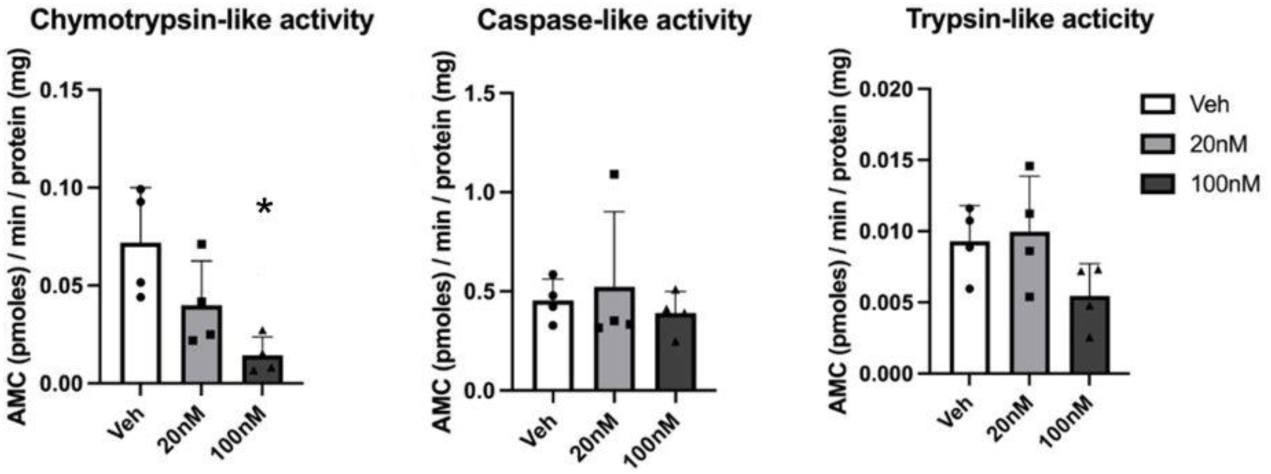
20S proteasome activities (chymotrypsin-like, caspase-like and trypsin-like activity from cortical cultures at 14DIV treated with veh (distilled water), 20nM or 100nM CORT. n=4/group. *p<0.05 vs vehicle group (post-hoc Tukey test). Results are shown as mean +/− s.e.m. with individual data points.

While PNNs are not detectable at 7 DIV, the widespread and diverse suppressive effects of GCs on the genes encoding components of PNNs that we have observed at 14 DIV strongly suggest that GC exposure is likely to have detrimental effects on PNN formation at this stage, but potentially not at 21 DIV. To clarify the impact of glucocorticoids on dendritic and structural formation of PNNs, WFA-labelling was monitored in cultured cells exposed to the same treatments.

At 14 DIV, exposure to 20nM or 100nM CORT resulted in decreased length of PNNs covering dendrites after 4h (F (2, 946) =56.11, p<0.001) (data not shown), with reduction of mean length from 29.3µm to 23.4µm with 20nM CORT (Tukey post hoc tests, veh vs 20nM CORT, p<0.001) and to 21.7µm with 100nM CORT exposure (Tukey post hoc tests, veh vs 100nM CORT, p<0.001). Length of dendrites covered by PNNs also decreased after 24h (F (2, 673) =14.89, p<0.001) CORT exposure (Fig.10 A,B), with reduction of mean length from 25.3µm to 21.0µm with 20nM CORT (Tukey post hoc tests, p<0.001) and to 23.4µm with 100nM CORT exposure (Tukey post hoc test, p=0.044). Similarly, the brightness intensity of WFA-labelled PNNs covering dendrites also decreased after 24h (F (2, 673) =14.89, p<0.001) with both 20nM and 100nM CORT treatment, from 23.09 a.u. to 20.13 a.u. and 16.9 a.u., respectively (p=0.024 Veh vs 20nM Tukey post hoc test; p<0.001, Veh vs high dose, Tukey post hoc test) (Fig.10 A,C); however, no significant changes in brightness of staining were found after 4h (F (2, 946 =1.12, p=0.327) treatment (data not shown).

**Figure 10.**
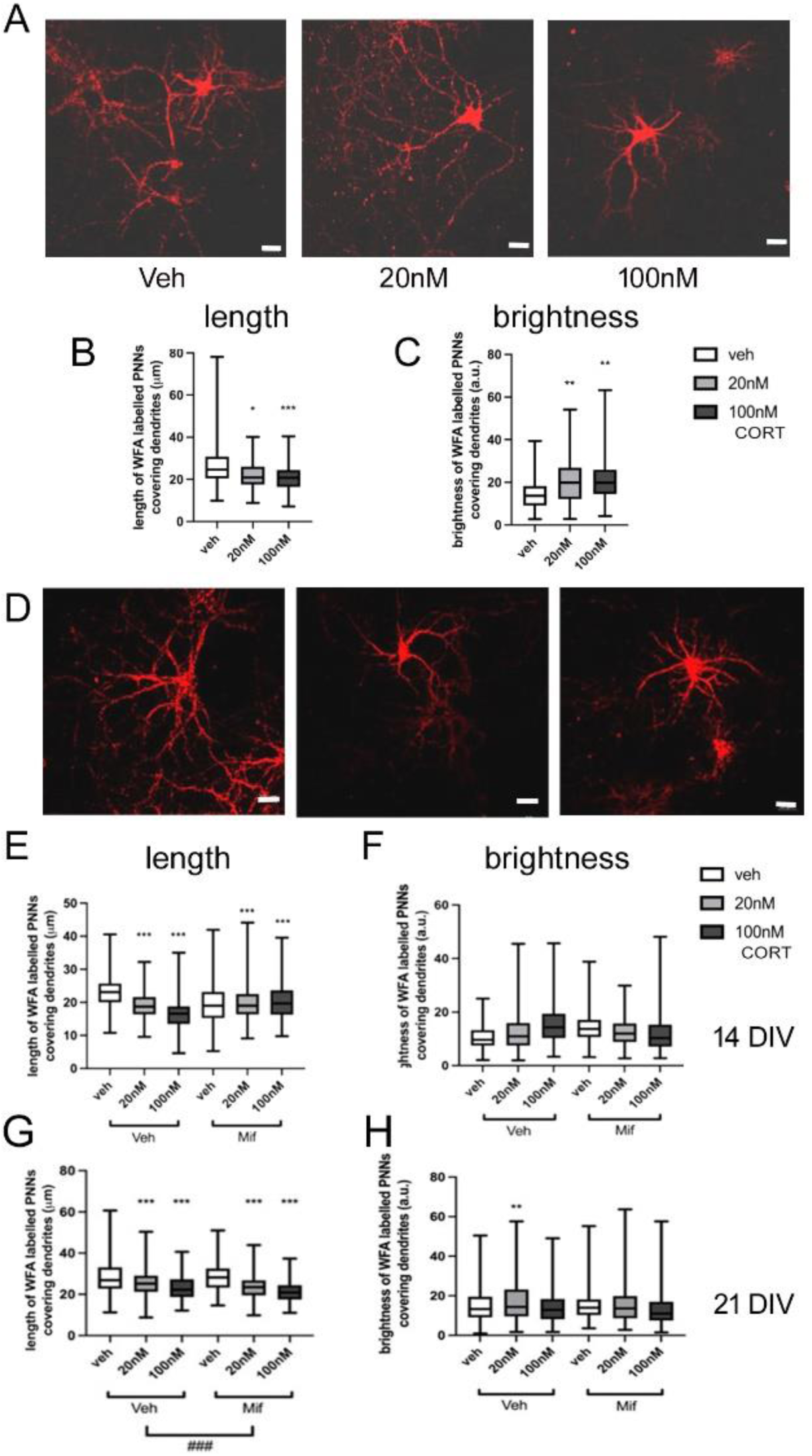
Effect of 24h glucocorticoid exposure on WFA-labelled PNNs in cultured neurons at 14 or 21 DIV **A**: representative images of WFA-labelled PNNs around cultured neurons at 14 DIV after 24h exposure to vehicle, 20nM or 100nM CORT. **B**: length of dendrite covered by PNN, and **C**: brightness intensity of WFA staining (Veh: 270 dendrites in 20 cells nested in 3 different slides with 2 cultures, 20nM: 166 dendrites in 20 cells nested in 3 different slides with 2 cultures, 100nM: 199 dendrites in 16 cells nested in 3 different slides with 2 cultures). **D**: representative images of WFA-labelled PNNs around cultured neurons at 14 DIV after 24h exposure to vehicle, 20nM or 100nM CORT in the presence of mifepristone (20nM). **E**: length of dendrite covered by PNN, and **F**: brightness intensity of WFA staining (Veh: veh: 139 dendrites in 13 cells nested in 3 different slides with 2 cultures, 20nM: 275 dendrites in 17 cells nested in 3 different slides with 2 cultures, 100nM: 164 dendrites in 15 cells nested in 3 different slides with 2 cultures; Mif: veh: 290 dendrites in 21 cells nested in 3 different slides with 2 cultures, 20nM: 182 dendrites in 16 cells nested in 3 different slides with 2 cultures, 100nM: 177 dendrites in 14 cells nested in 3 different slides with 2 cultures). **G,H:** Effects on PNNs at 21 DIV after 24 treatment **– G:** length of dendrite covered by PNN, and **H**: brightness intensity of WFA staining (Veh: veh: 360 dendrites in 32 cells nested in 3 different slides with 2 cultures, 20nM: 263 dendrites in 20 cells nested in 3 different slides with 2 cultures, 100nM: 265 dendrites in 20 cells nested in 3 different slides with 2 cultures; Mif: veh: 165 dendrites in 20 cells nested in 3 different slides with 2 cultures, 20nM: 297 dendrites in 18 cells nested in 3 different slides with 2 cultures, 100nM: 291 dendrites in 19 cells nested in 3 different slides with 2 cultures). ### p<0.001 ANOVA main effect mifepristone vs vehicle group ** p<0.01, *** p<0.001 post-hoc Tukey test vs corresponding vehicle control group. Scale bars represent 20μm. Boxes show median and interquartile range, with whiskers from minimum to maximum.

Moreover, at 21DIV, there was still an overall effect of CORT and of mifepristone on the length of PNN-covered dendrites after 24h treatment (F (1, 1580)) = 47.13, p<0.001). The interaction of CORT treatment and mifepristone was not quite significant (F (2, 1580)) = 2.81, p=0.060), but with a decreasing tendency in mean length of PNNs covering dendrites from 22.5µm to 19.0µm after 24h 100nM CORT with co-treated mifepristone (Tukey post hoc tests, p<0.001). However, no overall effect of mifepristone on the brightness intensity of WFA-labelled PNNs covered dendrites was found after 24h CORT and mifepristone treatment (F (1, 1580)) = 47.13, p=0.576) (fig.4 C D), however, there was a marginally significant interaction of CORT and mifepristone treatment (F (2, 1580)) = 3.06, p=0.047) with slightly increased brightness intensity of WFA-labelled PNNs after 100nM CORT and mifepristone treatment (Tukey post hoc test, p=0.046).

Hence we observe robust decreases in the length of proximal dendrite covered by PNN after CORT exposure, at both 14 and 21 DIV. These effects on PNN structure do not appear to be mediated via GRs.

Since loss of TnR reportedly reduces the number of dendrites that are covered proximally by PNNs (Weber et al., 1999), we also checked this parameter. However, no change in the number of WFA-labelled dendrites/cell was observed after CORT or mifepristone exposure (Supplementary figure S10).

## Discussion

A strength of this study is that the expression of all the genes encoding core PNN components has been monitored in parallel. Hence the relative effects on the different genes can be interpreted with confidence. Our initial expectation was that we might identify one or two components of the PNN where gene expression is modulated by CORT levels. Instead, we found a widespread and complex regulation of multiple PNN component genes, via diverse mechanisms, and suggesting a prominent role for GCs in modulating PNN gene expression, but potentially limited to early in development. The diminishing evidence for GC regulation of these genes from 7 to 21 DIV suggests that these gene expression mechanisms are very important while PNNs are forming and stabilising, but are much less important once they have attained a mature configuration.

Only for the phosphacan/*Ptprz1* gene was there a complete lack of evidence for regulation by GCs. Even in this case, GC modulation may be just hidden rather than absent. The exons encoding the phosphacan protein are also present in longer mRNAs encoding the full length Ptprz1 protein, and so any regulation specific for the phosphacan isoform might not be evident among the total transcripts, although a genomic action should still be evident, as, occurring prior to mRNA splicing, it should affect all protein isoforms.

### Mechanisms of GC regulation of PNN gene expression

Basal human serum cortisol concentrations are in the range of 150 to 350 nM, although around 80% is not free in solution, but is bound to cortisol-binding globulin (Hägg et al., 1987; Khalili-Mahani et al., 2015; Oster et al., 2016; Taylor et al., 1983), leaving free concentrations of around 30-70nM, although there is also marked circadian oscillation. Under conditions of high psychological stress, levels can rise to over 1000nM (Chatterton et al., 1997; Kotozaki & Kawashima, 2012; Vanhorebeek et al., 2006). Similarly, in mice, resting total corticosterone levels are around 100-350nM (free concentrations ∼ 20-70nM), rising to 1000-3000nM (free concentrations ∼ 200-600nM) during stress (Gong et al., 2015). During pregnancy, where mothers are not stressed, foetal cortisol levels have been estimated to be around 55nM (Gitau et al., 1998). Neuronal survival in culture relies on the presence of basal levels of GCs. Neurobasal medium with B27 supplement (which does not contain cortisol binding globulin) contains CORT concentrations of around 50-70nM (Brewer et al., 1993; Chen et al., 2008; Crochemore et al., 2005; Roth et al., 2010; Sünwoldt et al., 2017), in fact very similar to basal free GC concentrations in a non-stressed condition. Accordingly, we selected 2 concentrations of CORT to be added as experimental manipulation in this study, to represent both physiological (20nM) and pathological (100nM) levels of stress response. Resting GC concentrations will be fully activating MRs, whereas increasing concentrations under stress will additionally stimulate increasing proportions of GRs (and both the 20nM and 100nM concentrations will also activate GPR56/97 (Barros-Álvarez et al., 2022; Cain et al., 2023; de Kloet, 2022; Y. Q. Ping et al., 2021; Rafestin-Oblin et al., 1986).

In fact, where we observed effects of 100nM CORT, we generally also observed the effect with the lower concentration. At 7DIV, these effects were mostly mediated by GRs, as indicated by sensitivity to mifepristone. Where a significant effect was detected with the higher but not the lower concentration (as in suppression of *Gad1* and *Has3* expression), the same trend was observed with the lower concentration, consistent with dose-dependency at a single receptor. These effects at 7DIV, affecting many PNN component genes, reveal a widespread sensitivity of PNN component gene expression to GC levels at this developmental time, mediated by GRs.

At 7DIV, CORT suppressed *Ncan* and *TnR* expression, and mifepristone enhanced *Acan* expression and further suppressed *Ncan* and *TnR* expression. Thus basal GC levels are tending to promote *Ncan* and *TnR* expression relative to *Acan* expression, and there are additional non-GR-mediated GC actions to dampen down *Ncan* and *TnR* expression. Modulation of *Has* gene expression is prominent at this developmental stage - CORT suppressed *Has* 1,2 and 3 expression via GRs, whereas at 14DIV, this effect was not evident, but mifepristone elevated *Has* 1 and 2 expression. This implies that GCs are still acting to reduce *Has* 1 and 2 expression through GR activation at 14DIV, but that the effect has become more sensitive, so that the low GC concentrations in the culture medium are now effective. A powerful GC/GR-mediated suppression of *Has* 1,2 and 3 mRNA has been noted in peripheral cells (McRae et al., 2017; Stuhlmeier & Pollaschek, 2004; Zhang et al., 2000).

Apart from the elevated levels of *Has1,2* mRNAs induced by mifepristone at 14 DIV, decreased *Bcan* expression was also observed, despite the lack of GC modulation at 7 DIV, and suggesting heightened mRNA levels at 14 DIV due to basal GC levels. The other effect detected at 14DIV was a pronounced suppression of *Vcan* and *Hapln4* expression by CORT, where there also appeared to be increasing sensitivity to GCs, as no significant suppression was detected at 7 DIV, although, for the effect on *Vcan* mRNA levels at 14 DIV, GRs appeared not to be involved. Conversely, mRNA levels for *Ncan, TnR* and *Gad1* had now become insensitive to GCs.

While reduced expression of *Bcan, Ncan* and *Vcan* mRNA expression following GC exposure appears not to have been previously reported in the CNS, both the *Acan* and *Ncan* gene promoters contain GR-response elements (Rauch et al., 1995; Watanabe et al., 1995), and GC suppression of *Vcan* and *Acan* expression has been noted in peripheral cells (McRae et al., 2017; Short et al., 2020; Song et al., 2012). Interestingly, prenatal exposure to GCs downregulates peripheral tissue *Acan* expression (Chen et al., 2018), suggested that early developmental exposure can have lasting effects in offspring.

There is some existing evidence suggesting suppression of *Ncan* expression by stress or GC exposure. Liu et al. (Liu et al., 2008) reported GC-induced downregulation of expression *of Ncan* in astrocyte cultures (1 DIV) over 48h, partially mifepristone-sensitive, and intracerebral dexamethasone also reportedly decreased *Ncan* immunoreactivity (Zhong & Bellamkonda, 2007). Chronic stress in adolescence or in adulthood in rodents also seems to suppress cortical or hippocampal *Ncan* expression (Koskinen et al., 2020; Yu et al., 2022). Adult rodents exposed to stress show down-regulation at the protein level of hippocampal Bcan, Ncan, Ptprz1, TnR and Hapln1 but not Acan (Koskinen et al., 2020) and PFC Acan but not Bcan (Li et al., 2024).

The suppression of *Ncan* expression by GCs at 7 DIV was not attenuated by mifepristone, but a similar suppression was detected following exposure to collagen 3, an agonist at Gpr56 and Gpr97. It is interesting to note that *Ncan* appears to be synthesised primarily by astrocytes and glutamatergic projection neurons (Huntley et al., 2020; Irala et al., 2024; Zhang et al., 2014) (there will be a small proportion of astrocytes present in our neuronal-enriched cultures), and Gpr56 is expressed predominantly in astrocytes (but not glutamatergic projection neurons (Chiou et al., 2021; Huntley et al., 2020; Zhang et al., 2014). Hence it is possible that GCs can act via GRs to produce a generalised enhancement of Ncan expression, whereas GC activation of Gpr56 can reduce *Ncan* expression specifically in astrocytes.

GCs are reported to exert a post-transcriptional regulation of gene expression by GC/GR complexes affecting mRNA stability (Cho et al., 2015). However, we did not obtain any evidence to support this mechanism of action for the regulation of PNN component gene expression. Indeed, for many GC-induced changes in PNN component (suppression of *Bcan, Vcan* and *TnR* mRNA levels), we were unable to pinpoint the mechanisms of GC action. GRs appeared not to be involved (since mifepristone had little ability to attenuate the effects), and neither did Gpr56/97 (as reflected in the inability of collagen 3 to replicate the effect), but the effects seemed rapid, in that they were detectable within 4h, and so are likely to be non-genomic in nature.

### Effect on PNN structure

Compromised PNN formation (brightness of WFA staining) in cortex and hippocampus after exposure to prenatal or neonatal stress is well-documented in rats and mice (Allgäuer et al., 2023; Gildawie et al., 2020; Jakovljevic et al., 2022; Riga et al., 2017; Santiago et al., 2018). Most studies using mouse hippocampal or cortical cultures concur that at 14 DIV, PNNs are almost mature, with a net-like structure covering somata and proximal dendrites, and from then on the morphology changes little (Dityatev et al., 2007; Fowke et al., 2017; Geissler et al., 2013). Our observations were similar. This report may be the first demonstration that GCs modify PNN structure directly, and hence may be the mediators of the effects of stress on PNNs.

GC exposure caused a small but robust reduction in the length of PNN ensheathing the proximal dendrites, not involving GRs, and a more slowly-developing increase in intensity of WFA staining, likely to be mediated by GRs. The different mechanisms involved suggests that the increased brightness of WFA staining is not simply a result of compaction of a constant amount of PNN into a smaller volume. At 14 DIV, the magnitude of the changes in PNN structure might be considered smaller than expected, considering the quite profound changes in gene expression observed with GC exposure. However, PNNs seem to be quite robust to altered expression of component genes. While loss of Acan, phosphacan, TnR or Hapln4 compromises PNN structure (Bekku et al., 2012; Brückner et al., 2000; Eill et al., 2020; Giamanco et al., 2010; Haunsø et al., 2000; Morawski et al., 2014; Rowlands et al., 2018; Sucha et al., 2020; Suttkus et al., 2014; Weber et al., 1999), PNNs appear unaffected even in the complete absence of Bcan, Ncan or Has3 (Arranz et al., 2014; Brakebusch et al., 2002; Zhou et al., 2001).

The suppression of *Bcan*, *Vcan* and *Hapln4* gene expression by GCs at 14 DIV was not mifepristone-sensitive, and equally the decrement in PNN structure was also not mifepristone-sensitive. However it seems unlikely that the altered WFA staining is due to alterations in PNN component gene expression. Changes in PNN component gene expression detected when HCA was applied at 14 DIV were not attenuated by mifepristone, whereas alterations in WFA staining characteristics at 14 DIV were sensitive to mifepristone. Equally, no changes in PNN component gene expression were detected from GC exposure at 21 DIV, yet altered WFA staining characteristics are still observed. We have not systematically profiled PNN component protein expression at 21 DIV, so post-transcriptional actions of GCs may be responsible. For example, it has been suggested that Acan levels are partly controlled by post-translational modifications (Zhu, Cui, et al., 2019), so if these processes were modulated by GCs, a subtle but rapid effect on PNN structure might be evident.

### Modulation of GABAergic interneuron genes

Expression of both *Has3* and *Hapln4* (suppressed by GCs at 7 and 14 DIV) appears to be markedly enriched in Pvalb+ve cells as compared to other cortical neurons (Huntley et al., 2020). The regulation of *Has3* and *Hapln4* mRNA levels (and protein levels for Has3) by GCs might suggest a particular action of GCs on Pvalb+ve cells.

Elevated GC levels suppressed *Gad1* expression (7 and 14 DIV) and *Pvalb* expression (21 DIV), and at 7 DIV, blockade of GRs with mifepristone elevated expression of both *Gad1* and *Pvalb*. These 2 GABAergic interneuron genes, both of which show reduced expression levels in schizophrenia, are clearly sensitive to elevated GC levels. The action on *Gad1* expression at 7 DIV and 14 DIV appeared to involve GRs, whereas the action on *Pvalb* expression at 21 DIV, when PNN component genes were apparently resistant to GC actions, involved another mechanism.

These results are consistent with some previous observations. There are reports that chronic stress prenatally (Allgäuer et al., 2023; Heslin & Coutellier, 2018; Uchida et al., 2014; van de Looij et al., 2019), neonatally (Murthy et al., 2019) and in adult rodents (Banasr et al., 2017; Hu et al., 2010) decreases cortical *Pvalb* expression, although increased expression after adult stress has also been reported (Page et al., 2019). High GC concentrations for 72h also reportedly suppress *Gad1* (but not *Gad2*) expression in 10 DIV cultured cortical neurons (Banasr et al., 2017), and prenatal GC exposure *in vivo* reduces offspring *Pvalb* expression in hippocampus (Zhang et al., 2023). Hence our data, along with previous reports, seems clear in identifying Pvalb+ve cells potentially as direct targets of GCs for inducing gene expression changes relevant to schizophrenia.

### Proteasome

Proteasome inhibition leads to elevated cellular levels of proteins which are proteasome substrates. The modulation of neuronal proteasome activity by GCs has not previously been reported. There are some hints of proteasome modulation by GCs from peripheral cells, although for stimulation rather than inhibition. In hepatocytes, dexamethasone stimulates protein degradation within 4h (Hopgood et al, 1980), and similarly in muscle cells GCs accelerate protein degradation via ubiquitin pathways (Sun et al., 2008). There is a single report, in thymocytes, of physiologically-relevant concentrations of dexamethasone suppressing chymotrypsin-like and caspase-like, but not trypsin-like, proteasome activity within 3h (Beyette et al., 1998).

Here we report an inhibition of neuronal proteasome activity that is specific for chymotrypsin-like activity as compared to caspase-like (post-glutamate peptide hydrolase-like) activity (trypsin-like activity was very low, and it might be more difficult to observe suppression of activity). However, it should be noted that this suppression of chymotrypsin-like activity by CORT was demonstrated at 14DIV, while at this stage CORT decreased HAS2 protein levels, rather than the increase detected at 7DIV. It seems likely that a similar action of glucocorticoids on the proteasome occurs at 7DIV, but additional experiments would be needed to formally demonstrate this, potentially implicating suppression of proteasome activity in the Has2 protein increase. Has2 protein has an extremely short half-life (∼15minutes), and there is some evidence suggesting that its (lack of) longevity might be controlled by the proteasome at least in peripheral cells (Karousou et al., 2010; Vigetti et al., 2012), so elevated protein levels 4h after exposure to GCs are consistent with reduced degradation via proteasome inhibition.

## Conclusions

Perinatal trauma, which increases offspring schizophrenia risk (Davies et al., 2020; Paquin et al., 2021), elevates neonatal cortisol levels (Chiș et al., 2017; Gardner et al., 2001; Hernandez-Andrade et al., 2005; Improda et al., 2023). At this period, corresponding to our 7-14 DIV cultured neurons in mice, as PNNs form and stabilise, cortical *Ncan* and *Vcan* expression is declining, while *TnR* and *Pvalb* expression is increasing, in mice (Brückner et al., 2000; Du et al., 1996; Fertuzinhos et al., 2014; Fuss et al., 1993; Milev et al., 1998; Nowicka et al., 2009), and in humans (Honig et al., 1996; Kang et al., 2011; Letinic & Kostovic, 1998; Rogers et al., 2018). Our data suggest that inappropriately elevated cortisol levels at this time will be sufficient to disrupt the expression of PNN component genes at a critical time for PNN formation. We propose this mechanism as potentially contributing to the abnormal properties of Pvalb+ve interneurons in schizophrenia.

## Notes

### Competing Interest Statement

The authors have declared no competing interest.

## References

Allgäuer, L., Cabungcal, J. H., Yzydorczyk, C., Do, K. Q., & Dwir, D. (2023). Low protein-induced intrauterine growth restriction as a risk factor for schizophrenia phenotype in a rat model: assessing the role of oxidative stress and neuroinflammation interaction. Transl Psychiatry, 13(1), 30. 10.1038/s41398-023-02322-8

Arranz, A. M., Perkins, K. L., Irie, F., Lewis, D. P., Hrabe, J., Xiao, F., Itano, N., Kimata, K., Hrabetova, S., & Yamaguchi, Y. (2014). Hyaluronan deficiency due to Has3 knock-out causes altered neuronal activity and seizures via reduction in brain extracellular space. J Neurosci, 34(18), 6164–6176. 10.1523/jneurosci.3458-13.2014

Banasr, M., Lepack, A., Fee, C., Duric, V., Maldonado-Aviles, J., DiLeone, R., Sibille, E., Duman, R. S., & Sanacora, G. (2017). Characterization of GABAergic marker expression in the chronic unpredictable stress model of depression. Chronic Stress (Thousand Oaks), 1. 10.1177/2470547017720459

Barros-Álvarez, X., Nwokonko, R. M., Vizurraga, A., Matzov, D., He, F., Papasergi-Scott, M. M., Robertson, M. J., Panova, O., Yardeni, E. H., Seven, A. B., Kwarcinski, F. E., Su, H., Peroto, M. C., Meyerowitz, J. G., Shalev-Benami, M., Tall, G. G., & Skiniotis, G. (2022). The tethered peptide activation mechanism of adhesion GPCRs. Nature, 604(7907), 757–762. 10.1038/s41586-022-04575-7

Bekku, Y., Saito, M., Moser, M., Fuchigami, M., Maehara, A., Nakayama, M., Kusachi, S., Ninomiya, Y., & Oohashi, T. (2012). Bral2 is indispensable for the proper localization of brevican and the structural integrity of the perineuronal net in the brainstem and cerebellum. J Comp Neurol, 520(8), 1721–1736. 10.1002/cne.23009

Beyette, J., Mason, G. G., Murray, R. Z., Cohen, G. M., & Rivett, A. J. (1998). Proteasome activities decrease during dexamethasone-induced apoptosis of thymocytes. Biochem J, 332 (Pt 2)(Pt 2), 315–320. 10.1042/bj3320315

Brakebusch, C., Seidenbecher, C. I., Asztely, F., Rauch, U., Matthies, H., Meyer, H., Krug, M., Böckers, T. M., Zhou, X., Kreutz, M. R., Montag, D., Gundelfinger, E. D., & Fässler, R. (2002). Brevican-deficient mice display impaired hippocampal CA1 long-term potentiation but show no obvious deficits in learning and memory. Mol Cell Biol, 22(21), 7417–7427. 10.1128/mcb.22.21.7417-7427.2002

Brewer, G. J., Torricelli, J. R., Evege, E. K., & Price, P. J. (1993). Optimized survival of hippocampal neurons in B27-supplemented Neurobasal, a new serum-free medium combination. J Neurosci Res, 35(5), 567–576. 10.1002/jnr.490350513

Brückner, G., Grosche, J., Schmidt, S., Härtig, W., Margolis, R. U., Delpech, B., Seidenbecher, C. I., Czaniera, R., & Schachner, M. (2000). Postnatal development of perineuronal nets in wild-type mice and in a mutant deficient in tenascin-R. J Comp Neurol, 428(4), 616–629. 10.1002/1096-9861(20001225)428:4<616::aid-cne3>3.0.co;2-k

Cain, D., Cidlowski, J., Edwards, D. P., Fuller, P., Grimm, S. L., Hartig, S., Lange, C. A., Oakley, R. H., Richer, J. K., Sartorius, C. A., Tetel, M., Weigel, N., & Young, M. J. (2023). 3C. 3-Ketosteroid receptors in GtoPdb v.2023.1. IUPHAR/BPS Guide to Pharmacology CITE, 2023(1). 10.2218/gtopdb/F98/2023.1

Celio, M. R., & Chiquet-Ehrismann, R. (1993). ’Perineuronal nets’ around cortical interneurons expressing parvalbumin are rich in tenascin. Neurosci Lett, 162(1-2), 137–140. 10.1016/0304-3940(93)90579-a

Chatterton, R. T., Jr., Vogelsong, K. M., Lu, Y.-c., & Hudgens, G. A. (1997). Hormonal Responses to Psychological Stress in Men Preparing for Skydiving1. The Journal of Clinical Endocrinology & Metabolism, 82(8), 2503–2509. 10.1210/jcem.82.8.4133

Chen, Y., Stevens, B., Chang, J., Milbrandt, J., Barres, B. A., & Hell, J. W. (2008). NS21: re-defined and modified supplement B27 for neuronal cultures. J Neurosci Methods, 171(2), 239–247. 10.1016/j.jneumeth.2008.03.013

Chen, Z., Zhao, Z., Li, Y., Zhang, X., Li, B., Chen, L., & Wang, H. (2018). Course-, dose-, and stage-dependent toxic effects of prenatal dexamethasone exposure on fetal articular cartilage development. Toxicology Letters, 286, 1–9. 10.1016/j.toxlet.2018.01.008

Chiou, B., Gao, C., Giera, S., Folts, C. J., Kishore, P., Yu, D., Oak, H. C., Jiang, R., & Piao, X. (2021). Cell type-specific evaluation of ADGRG1/GPR56 function in developmental central nervous system myelination. Glia, 69(2), 413–423. 10.1002/glia.23906

Chiș, A., Vulturar, R., Andreica, S., Prodan, A., & Miu, A. C. (2017). Behavioral and cortisol responses to stress in newborn infants: Effects of mode of delivery. Psychoneuroendocrinology, 86, 203–208. 10.1016/j.psyneuen.2017.09.024

Crochemore, C., Lu, J., Wu, Y., Liposits, Z., Sousa, N., Holsboer, F., & Almeida, O. F. X. (2005). Direct targeting of hippocampal neurons for apoptosis by glucocorticoids is reversible by mineralocorticoid receptor activation. Molecular Psychiatry, 10(8), 790–798. 10.1038/sj.mp.4001679

Cruise, L., Ho, L. K., Veitch, K., Fuller, G., & Morris, B. J. (2000). Kainate receptors activate NFkB via MAP kinase in striatal neurones. Neuroreport, 11, 395–401.

Davies, C., Segre, G., Estradé, A., Radua, J., De Micheli, A., Provenzani, U., Oliver, D., Salazar de Pablo, G., Ramella-Cravaro, V., Besozzi, M., Dazzan, P., Miele, M., Caputo, G., Spallarossa, C., Crossland, G., Ilyas, A., Spada, G., Politi, P., Murray, R. M., … Fusar-Poli, P. (2020). Prenatal and perinatal risk and protective factors for psychosis: a systematic review and meta-analysis. Lancet Psychiatry, 7(5), 399–410. 10.1016/s2215-0366(20)30057-2

De Kloet, E. R. (2004). Hormones and the stressed brain. Ann N Y Acad Sci, 1018, 1–15. 10.1196/annals.1296.001

de Kloet, E. R. (2022). Brain mineralocorticoid and glucocorticoid receptor balance in neuroendocrine regulation and stress-related psychiatric etiopathologies. Current Opinion in Endocrine and Metabolic Research, 24, 100352. 10.1016/j.coemr.2022.100352

Dityatev, A., Brückner, G., Dityateva, G., Grosche, J., Kleene, R., & Schachner, M. (2007). Activity-dependent formation and functions of chondroitin sulfate-rich extracellular matrix of perineuronal nets. Dev Neurobiol, 67(5), 570–588. 10.1002/dneu.20361

Du, J., Zhang, L., Weiser, M., Rudy, B., & McBain, C. (1996). Developmental expression and functional characterization of the potassium-channel subunit Kv3.1b in parvalbumin-containing interneurons of the rat hippocampus. The Journal of Neuroscience, 16(2), 506–518. 10.1523/jneurosci.16-02-00506.1996

Eill, G. J., Sinha, A., Morawski, M., Viapiano, M. S., & Matthews, R. T. (2020). The protein tyrosine phosphatase RPTPζ/phosphacan is critical for perineuronal net structure. J Biol Chem, 295(4), 955–968. 10.1074/jbc.RA119.010830

Enwright, J. F., Sanapala, S., Foglio, A., Berry, R., Fish, K. N., & Lewis, D. A. (2016). Reduced Labeling of Parvalbumin Neurons and Perineuronal Nets in the Dorsolateral Prefrontal Cortex of Subjects with Schizophrenia. neuropsychopharmacology, 41(9), 2206–2214. 10.1038/npp.2016.24

Fawcett, J. W., Oohashi, T., & Pizzorusso, T. (2019). The roles of perineuronal nets and the perinodal extracellular matrix in neuronal function. Nat Rev Neurosci, 20(8), 451–465. 10.1038/s41583-019-0196-3

Ferrer-Ferrer, M., & Dityatev, A. (2018). Shaping Synapses by the Neural Extracellular Matrix. Front Neuroanat, 12, 40. 10.3389/fnana.2018.00040

Fertuzinhos, S., Li, M., Kawasawa, Y. I., Ivic, V., Franjic, D., Singh, D., Crair, M., & Sestan, N. (2014). Laminar and temporal expression dynamics of coding and noncoding RNAs in the mouse neocortex. Cell Rep, 6(5), 938–950. 10.1016/j.celrep.2014.01.036

Fowden, A. L., Vaughan, O. R., Murray, A. J., & Forhead, A. J. (2022). Metabolic Consequences of Glucocorticoid Exposure before Birth. Nutrients, 14(11). 10.3390/nu14112304

Fowke, T. M., Karunasinghe, R. N., Bai, J. Z., Jordan, S., Gunn, A. J., & Dean, J. M. (2017). Hyaluronan synthesis by developing cortical neurons in vitro. Sci Rep, 7, 44135. 10.1038/srep44135

Fuller, G., Veitch, K., Lai Kwan, H., Cruise, L., & Morris, B. J. (2001). Activation of p44/p42 MAP kinase in striatal neurons via kainate receptors and PI3 kinase. Molecular Brain Research, 89(1-2), 126–132.

Fuss, B., Wintergerst, E. S., Bartsch, U., & Schachner, M. (1993). Molecular characterization and in situ mRNA localization of the neural recognition molecule J1-160/180: a modular structure similar to tenascin. J Cell Biol, 120(5), 1237–1249. 10.1083/jcb.120.5.1237

Gardner, D. S., Fletcher, A. J., Fowden, A. L., & Giussani, D. A. (2001). Plasma adrenocorticotropin and cortisol concentrations during acute hypoxemia after a reversible period of adverse intrauterine conditions in the ovine fetus during late gestation. Endocrinology, 142(2), 589–598. 10.1210/endo.142.2.7980

Geissler, M., Gottschling, C., Aguado, A., Rauch, U., Wetzel, C. H., Hatt, H., & Faissner, A. (2013). Primary hippocampal neurons, which lack four crucial extracellular matrix molecules, display abnormalities of synaptic structure and function and severe deficits in perineuronal net formation. J Neurosci, 33(18), 7742–7755. 10.1523/jneurosci.3275-12.2013

Giamanco, K. A., Morawski, M., & Matthews, R. T. (2010). Perineuronal net formation and structure in aggrecan knockout mice. Neuroscience, 170(4), 1314–1327. 10.1016/j.neuroscience.2010.08.032

Gildawie, K. R., Honeycutt, J. A., & Brenhouse, H. C. (2020). Region-specific Effects of Maternal Separation on Perineuronal Net and Parvalbumin-expressing Interneuron Formation in Male and Female Rats. Neuroscience, 428, 23–37. 10.1016/j.neuroscience.2019.12.010

Gitau, R., Cameron, A., Fisk, N. M., & Glover, V. (1998). Fetal exposure to maternal cortisol. lancet, 352(9129), 707–708. 10.1016/s0140-6736(05)60824-0

Gong, S., Miao, Y.-L., Jiao, G.-Z., Sun, M.-J., Li, H., Lin, J., Luo, M.-J., & Tan, J.-H. (2015). Dynamics and Correlation of Serum Cortisol and Corticosterone under Different Physiological or Stressful Conditions in Mice. PLoS ONE, 10(2), e0117503. 10.1371/journal.pone.0117503

Gonzalez-Burgos, G., Cho, R. Y., & Lewis, D. A. (2015). Alterations in cortical network oscillations and parvalbumin neurons in schizophrenia. Biological Psychiatry, 77(12), 1031–1040. 10.1016/j.biopsych.2015.03.010

Hägg, E., Asplund, K., & Lithner, F. (1987). Value of basal plasma cortisol assays in the assessment of pituitary-adrenal insufficiency. Clin Endocrinol (Oxf), 26(2), 221–226. 10.1111/j.1365-2265.1987.tb00780.x

Härtig, W., Brauer, K., & Brückner, G. (1992). Wisteria floribunda agglutinin-labelled nets surround parvalbumin-containing neurons. Neuroreport, 3(10), 869–872. 10.1097/00001756-199210000-00012

Haunsø, A., Ibrahim, M., Bartsch, U., Letiembre, M., Celio, M. R., & Menoud, P. (2000). Morphology of perineuronal nets in tenascin-R and parvalbumin single and double knockout mice. Brain Res, 864(1), 142–145. 10.1016/s0006-8993(00)02173-9

Hernandez-Andrade, E., Hellström-Westas, L., Thorngren-Jerneck, K., Jansson, T., Liuba, K., Lingman, G., Marsál, K., Oskarsson, G., Werner, O., & Ley, D. (2005). Perinatal adaptive response of the adrenal and carotid blood flow in sheep fetuses subjected to total cord occlusion. J Matern Fetal Neonatal Med, 17(2), 101–109. 10.1080/14767050500043509

Heslin, K., & Coutellier, L. (2018). Npas4 deficiency and prenatal stress interact to affect social recognition in mice. Genes Brain Behav, 17(5), e12448. 10.1111/gbb.12448

Hoftman, G. D., Volk, D. W., Bazmi, H. H., Li, S., Sampson, A. R., & Lewis, D. A. (2015). Altered cortical expression of GABA-related genes in schizophrenia: illness progression vs developmental disturbance. Schizophrenia Bulletin, 41(1), 180–191. 10.1093/schbul/sbt178

Hollos, P., John, J. M., Lehtonen, J. V., & Coffey, E. T. (2020). Optogenetic Control of Spine-Head JNK Reveals a Role in Dendritic Spine Regression. eNeuro, 7(1). 10.1523/eneuro.0303-19.2019

Honig, L. S., Herrmann, K., & Shatz, C. J. (1996). Developmental Changes Revealed by Immunohistochemical Markers in Human Cerebral Cortex. Cerebral Cortex, 6(6), 794–806. 10.1093/cercor/6.6.794

Hu, W., Zhang, M., Czéh, B., Flügge, G., & Zhang, W. (2010). Stress impairs GABAergic network function in the hippocampus by activating nongenomic glucocorticoid receptors and affecting the integrity of the parvalbumin-expressing neuronal network. neuropsychopharmacology, 35(8), 1693–1707. 10.1038/npp.2010.31

Huntley, M. A., Srinivasan, K., Friedman, B. A., Wang, T. M., Yee, A. X., Wang, Y., Kaminker, J. S., Sheng, M., Hansen, D. V., & Hanson, J. E. (2020). Genome-Wide Analysis of Differential Gene Expression and Splicing in Excitatory Neurons and Interneuron Subtypes. J Neurosci, 40(5), 958–973. 10.1523/jneurosci.1615-19.2019

Hynes, D., & Harvey, B. J. (2019). Dexamethasone reduces airway epithelial Cl(-) secretion by rapid non-genomic inhibition of KCNQ1, KCNN4 and KATP K(+) channels. Steroids, 151, 108459. 10.1016/j.steroids.2019.108459

Improda, N., Capalbo, D., Poloniato, A., Garbetta, G., Dituri, F., Penta, L., Aversa, T., Sessa, L., Vierucci, F., Cozzolino, M., Vigone, M. C., Tronconi, G. M., Del Pistoia, M., Lucaccioni, L., Tuli, G., Munarin, J., Tessaris, D., de Sanctis, L., & Salerno, M. (2023). Perinatal asphyxia and hypothermic treatment from the endocrine perspective. Front Endocrinol (Lausanne), 14, 1249700. 10.3389/fendo.2023.1249700

Irala, D., Wang, S., Sakers, K., Nagendren, L., Ulloa Severino, F. P., Bindu, D. S., Savage, J. T., & Eroglu, C. (2024). Astrocyte-secreted neurocan controls inhibitory synapse formation and function. Neuron, 112(10), 1657–1675.e1610. 10.1016/j.neuron.2024.03.007

Jakovljevic, A., Agatonovic, G., Aleksic, D., Aksic, M., Reiss, G., Förster, E., Stamatakis, A., Jakovcevski, I., & Poleksic, J. (2022). The impact of early life maternal deprivation on the perineuronal nets in the prefrontal cortex and hippocampus of young adult rats [Original Research]. Frontiers in Cell and Developmental Biology, 10. 10.3389/fcell.2022.982663

Joëls, M. (2018). Corticosteroids and the brain. J Endocrinol, 238(3), R121–r130. 10.1530/joe-18-0226

Joëls, M., Pasricha, N., & Karst, H. (2013). The interplay between rapid and slow corticosteroid actions in brain. Eur J Pharmacol, 719(1-3), 44–52. 10.1016/j.ejphar.2013.07.015

John, N., Krügel, H., Frischknecht, R., Smalla, K. H., Schultz, C., Kreutz, M. R., Gundelfinger, E. D., & Seidenbecher, C. I. (2006). Brevican-containing perineuronal nets of extracellular matrix in dissociated hippocampal primary cultures. molecular and cellular neurosciences, 31(4), 774–784. 10.1016/j.mcn.2006.01.011

Kang, H. J., Kawasawa, Y. I., Cheng, F., Zhu, Y., Xu, X., Li, M., Sousa, A. M., Pletikos, M., Meyer, K. A., Sedmak, G., Guennel, T., Shin, Y., Johnson, M. B., Krsnik, Z., Mayer, S., Fertuzinhos, S., Umlauf, S., Lisgo, S. N., Vortmeyer, A., … Sestan, N. (2011). Spatio-temporal transcriptome of the human brain. Nature, 478(7370), 483–489. 10.1038/nature10523

Karousou, E., Kamiryo, M., Skandalis, S. S., Ruusala, A., Asteriou, T., Passi, A., Yamashita, H., Hellman, U., Heldin, C. H., & Heldin, P. (2010). The activity of hyaluronan synthase 2 is regulated by dimerization and ubiquitination. J Biol Chem, 285(31), 23647–23654. 10.1074/jbc.M110.127050

Khalili-Mahani, N., Martini, C. H., Olofsen, E., Dahan, A., & Niesters, M. (2015). Effect of subanaesthetic ketamine on plasma and saliva cortisol secretion. British Journal of Anaesthesia, 115(1), 68–75. 10.1093/bja/aev135

Koskinen, M. K., van Mourik, Y., Smit, A. B., Riga, D., & Spijker, S. (2020). From stress to depression: development of extracellular matrix-dependent cognitive impairment following social stress. Sci Rep, 10(1), 17308. 10.1038/s41598-020-73173-2

Kotozaki, Y., & Kawashima, R. (2012). Effects of the Higashi-Nihon Earthquake: Posttraumatic Stress, Psychological Changes, and Cortisol Levels of Survivors. PLoS ONE, 7(4), e34612. 10.1371/journal.pone.0034612

Letinic, K., & Kostovic, I. (1998). Postnatal development of calcium-binding proteins calbindin and parvalbumin in human visual cortex. Cereb Cortex, 8(7), 660–669. 10.1093/cercor/8.7.660

Li, X., Ren, D., Luo, B., Liu, Z., Li, N., Zhou, T., & Fei, E. (2024). Perineuronal Nets Alterations Contribute to Stress-Induced Anxiety-Like Behavior. Mol Neurobiol, 61(1), 411–422. 10.1007/s12035-023-03596-1

Liu, W. L., Lee, Y. H., Tsai, S. Y., Hsu, C. Y., Sun, Y. Y., Yang, L. Y., Tsai, S. H., & Yang, W. C. (2008). Methylprednisolone inhibits the expression of glial fibrillary acidic protein and chondroitin sulfate proteoglycans in reactivated astrocytes. Glia, 56(13), 1390–1400. 10.1002/glia.20706

Luo, R., Jeong, S. J., Jin, Z., Strokes, N., Li, S., & Piao, X. (2011). G protein-coupled receptor 56 and collagen III, a receptor-ligand pair, regulates cortical development and lamination. Proc Natl Acad Sci U S A, 108(31), 12925–12930. 10.1073/pnas.1104821108

Mauney, S. A., Athanas, K. M., Pantazopoulos, H., Shaskan, N., Passeri, E., Berretta, S., & Woo, T. U. (2013). Developmental pattern of perineuronal nets in the human prefrontal cortex and their deficit in schizophrenia. Biological Psychiatry, 74(6), 427–435. 10.1016/j.biopsych.2013.05.007

Mawson, E. R., & Morris, B. J. (2023). A consideration of the increased risk of schizophrenia due to prenatal maternal stress, and the possible role of microglia. progress in neuro-psychopharmacology and biological psychiatry, 125, 110773. 10.1016/j.pnpbp.2023.110773

McNair, K., Spike, R., Guilding, C., Prendergast, G. C., Stone, T. W., Cobb, S. R., & Morris, B. J. (2010). A role for RhoB in synaptic plasticity and the regulation of neuronal morphology. J Neurosci, 30(9), 3508–3517. 10.1523/JNEUROSCI.5386-09.2010.

McRae, N., Forgan, L., McNeill, B., Addinsall, A., McCulloch, D., Van der Poel, C., & Stupka, N. (2017). Glucocorticoids Improve Myogenic Differentiation In Vitro by Suppressing the Synthesis of Versican, a Transitional Matrix Protein Overexpressed in Dystrophic Skeletal Muscles. Int J Mol Sci, 18(12). 10.3390/ijms18122629

Milev, P., Maurel, P., Chiba, A., Mevissen, M., Popp, S., Yamaguchi, Y., Margolis, R. K., & Margolis, R. U. (1998). Differential regulation of expression of hyaluronan-binding proteoglycans in developing brain: aggrecan, versican, neurocan, and brevican. Biochem Biophys Res Commun, 247(2), 207–212. 10.1006/bbrc.1998.8759

Morawski, M., Dityatev, A., Hartlage-Rübsamen, M., Blosa, M., Holzer, M., Flach, K., Pavlica, S., Dityateva, G., Grosche, J., Brückner, G., & Schachner, M. (2014). Tenascin-R promotes assembly of the extracellular matrix of perineuronal nets via clustering of aggrecan. Philos Trans R Soc Lond B Biol Sci, 369(1654), 20140046. 10.1098/rstb.2014.0046

Morawski, M., Reinert, T., Meyer-Klaucke, W., Wagner, F. E., Tröger, W., Reinert, A., Jäger, C., Brückner, G., & Arendt, T. (2015). Ion exchanger in the brain: Quantitative analysis of perineuronally fixed anionic binding sites suggests diffusion barriers with ion sorting properties. Sci Rep, 5, 16471. 10.1038/srep16471

Morishita, H., Cabungcal, J. H., Chen, Y., Do, K. Q., & Hensch, T. K. (2015). Prolonged Period of Cortical Plasticity upon Redox Dysregulation in Fast-Spiking Interneurons. Biological Psychiatry, 78(6), 396–402. 10.1016/j.biopsych.2014.12.026

Murthy, S., Kane, G. A., Katchur, N. J., Lara Mejia, P. S., Obiofuma, G., Buschman, T. J., McEwen, B. S., & Gould, E. (2019). Perineuronal Nets, Inhibitory Interneurons, and Anxiety-Related Ventral Hippocampal Neuronal Oscillations Are Altered by Early Life Adversity. Biological Psychiatry, 85(12), 1011–1020. 10.1016/j.biopsych.2019.02.021

Nowicka, D., Soulsby, S., Skangiel-Kramska, J., & Glazewski, S. (2009). Parvalbumin-containing neurons, perineuronal nets and experience-dependent plasticity in murine barrel cortex. Eur J Neurosci, 30(11), 2053–2063. 10.1111/j.1460-9568.2009.06996.x

O’Kane, E. M., Stone, T. W., & Morris, B. J. (2003). Activation of Rho GTPases by synaptic transmission in the hippocampus. Journal of Neurochemistry, 87, 1309–1312.

Olaniru, O. E., Pingitore, A., Giera, S., Piao, X., Castañera González, R., Jones, P. M., & Persaud, S. J. (2018). The adhesion receptor GPR56 is activated by extracellular matrix collagen III to improve β-cell function. Cellular and molecular life sciences : CMLS, 75(21), 4007–4019. 10.1007/s00018-018-2846-4

Oster, H., Challet, E., Ott, V., Arvat, E., de Kloet, E. R., Dijk, D.-J., Lightman, S., Vgontzas, A., & Van Cauter, E. (2016). The Functional and Clinical Significance of the 24-Hour Rhythm of Circulating Glucocorticoids. Endocrine Reviews, 38(1), 3–45. 10.1210/er.2015-1080

Page, C. E., Shepard, R., Heslin, K., & Coutellier, L. (2019). Prefrontal parvalbumin cells are sensitive to stress and mediate anxiety-related behaviors in female mice. Sci Rep, 9(1), 19772. 10.1038/s41598-019-56424-9

Pantazopoulos, H., Katsel, P., Haroutunian, V., Chelini, G., Klengel, T., & Berretta, S. (2021). Molecular signature of extracellular matrix pathology in schizophrenia. Eur J Neurosci, 53(12), 3960–3987. 10.1111/ejn.15009

Paquin, V., Lapierre, M., Veru, F., & King, S. (2021). Early Environmental Upheaval and the Risk for Schizophrenia. Annu Rev Clin Psychol, 17, 285–311. 10.1146/annurev-clinpsy-081219-103805

Paterson, G. J., Ohashi, Y., Reynolds, G. P., Pratt, J., & Morris, B. J. (2006). Selective increases in the cytokine, TNFα, in the prefrontal cortex of PCP-treated rats and human schizophrenic subjects: influence of antipsychotic drugs. J Psychopharmacol, 20, 636–642. http://jop.sagepub.com/cgi/content/abstract/0269881106062025v1

Ping, Y.-Q., Mao, C., Xiao, P., Zhao, R.-J., Jiang, Y., Yang, Z., An, W.-T., Shen, D.-D., Yang, F., Zhang, H., Qu, C., Shen, Q., Tian, C., Li, Z.-j., Li, S., Wang, G.-Y., Tao, X., Wen, X., Zhong, Y.-N., … Sun, J.-P. (2021). Structures of the glucocorticoid-bound adhesion receptor GPR97–Go complex. Nature, 589(7843), 620–626. 10.1038/s41586-020-03083-w

Ping, Y. Q., Mao, C., Xiao, P., Zhao, R. J., Jiang, Y., Yang, Z., An, W. T., Shen, D. D., Yang, F., Zhang, H., Qu, C., Shen, Q., Tian, C., Li, Z. J., Li, S., Wang, G. Y., Tao, X., Wen, X., Zhong, Y. N., … Sun, J. P. (2021). Structures of the glucocorticoid-bound adhesion receptor GPR97-G(o) complex. Nature, 589(7843), 620–626. 10.1038/s41586-020-03083-w

Pizzorusso, T., Medini, P., Berardi, N., Chierzi, S., Fawcett, J. W., & Maffei, L. (2002). Reactivation of ocular dominance plasticity in the adult visual cortex. Science, 298(5596), 1248–1251. 10.1126/science.1072699

Rafestin-Oblin, M. E., Lombes, M., Lustenberger, P., Blanchardie, P., Michaud, A., Cornu, G., & Claire, M. (1986). Affinity of corticosteroids for mineralocorticoid and glucocorticoid receptors of the rabbit kidney: effect of steroid substitution. J Steroid Biochem, 25(4), 527–534. 10.1016/0022-4731(86)90398-5

Rauch, U., Grimpe, B., Kulbe, G., Arnold-Ammer, I., Beier, D. R., & Fässler, R. (1995). Structure and chromosomal localization of the mouse neurocan gene. Genomics, 28(3), 405–410. 10.1006/geno.1995.1168

Riga, D., Kramvis, I., Koskinen, M. K., van Bokhoven, P., van der Harst, J. E., Heistek, T. S., Jaap Timmerman, A., van Nierop, P., van der Schors, R. C., Pieneman, A. W., de Weger, A., van Mourik, Y., Schoffelmeer, A. N. M., Mansvelder, H. D., Meredith, R. M., Hoogendijk, W. J. G., Smit, A. B., & Spijker, S. (2017). Hippocampal extracellular matrix alterations contribute to cognitive impairment associated with a chronic depressive-like state in rats. Sci Transl Med, 9(421). 10.1126/scitranslmed.aai8753

Rogers, S. L., Rankin-Gee, E., Risbud, R. M., Porter, B. E., & Marsh, E. D. (2018). Normal Development of the Perineuronal Net in Humans; In Patients with and without Epilepsy. Neuroscience, 384, 350–360. 10.1016/j.neuroscience.2018.05.039

Roth, S., Zhang, S., Chiu, J., Wirth, E. K., & Schweizer, U. (2010). Development of a serum-free supplement for primary neuron culture reveals the interplay of selenium and vitamin E in neuronal survival. J Trace Elem Med Biol, 24(2), 130–137. 10.1016/j.jtemb.2010.01.007

Rowlands, D., Lensjø, K. K., Dinh, T., Yang, S., Andrews, M. R., Hafting, T., Fyhn, M., Fawcett, J. W., & Dick, G. (2018). Aggrecan Directs Extracellular Matrix-Mediated Neuronal Plasticity. J Neurosci, 38(47), 10102–10113. 10.1523/jneurosci.1122-18.2018

Santiago, A. N., Lim, K. Y., Opendak, M., Sullivan, R. M., & Aoki, C. (2018). Early life trauma increases threat response of peri-weaning rats, reduction of axo-somatic synapses formed by parvalbumin cells and perineuronal net in the basolateral nucleus of amygdala. J Comp Neurol, 526(16), 2647–2664. 10.1002/cne.24522

Schneider, C. A., Rasband, W. S., & Eliceiri, K. W. (2012). NIH Image to ImageJ: 25 years of image analysis. Nature methods, 9(7), 671–675. 10.1038/nmeth.2089

Schwartz, L. B. (1997). Understanding human parturition. lancet, 350(9094), 1792–1793. 10.1016/s0140-6736(05)63632-x

Short, K. L., Bird, A. D., Seow, B. K. L., Ng, J., McDougall, A. R. A., Wallace, M. J., Hooper, S. B., & Cole, T. J. (2020). Glucocorticoid signalling drives reduced versican levels in the fetal mouse lung. J Mol Endocrinol, 64(3), 155–164. 10.1530/jme-19-0235

Song, Y. W., Zhang, T., & Wang, W. B. (2012). Gluococorticoid could influence extracellular matrix synthesis through Sox9 via p38 MAPK pathway. Rheumatol Int, 32(11), 3669–3673. 10.1007/s00296-011-2091-8

Steullet, P., Cabungcal, J. H., Bukhari, S. A., Ardelt, M. I., Pantazopoulos, H., Hamati, F., Salt, T. E., Cuenod, M., Do, K. Q., & Berretta, S. (2018). The thalamic reticular nucleus in schizophrenia and bipolar disorder: role of parvalbumin-expressing neuron networks and oxidative stress. Molecular Psychiatry, 23(10), 2057–2065. 10.1038/mp.2017.230

Stuhlmeier, K. M., & Pollaschek, C. (2004). Glucocorticoids inhibit induced and non-induced mRNA accumulation of genes encoding hyaluronan synthases (HAS): hydrocortisone inhibits HAS1 activation by blocking the p38 mitogen-activated protein kinase signalling pathway. Rheumatology (Oxford), 43(2), 164–169. 10.1093/rheumatology/keh014

Sucha, P., Chmelova, M., Kamenicka, M., Bochin, M., Oohashi, T., & Vargova, L. (2020). The Effect of Hapln4 Link Protein Deficiency on Extracellular Space Diffusion Parameters and Perineuronal Nets in the Auditory System During Aging. Neurochem Res, 45(1), 68–82. 10.1007/s11064-019-02894-2

Sun, L., Trausch-Azar, J. S., Muglia, L. J., & Schwartz, A. L. (2008). Glucocorticoids differentially regulate degradation of MyoD and Id1 by N-terminal ubiquitination to promote muscle protein catabolism. Proc Natl Acad Sci U S A, 105(9), 3339–3344. 10.1073/pnas.0800165105

Sünwoldt, J., Bosche, B., Meisel, A., & Mergenthaler, P. (2017). Neuronal Culture Microenvironments Determine Preferences in Bioenergetic Pathway Use. Front Mol Neurosci, 10, 305. 10.3389/fnmol.2017.00305

Suttkus, A., Rohn, S., Weigel, S., Glöckner, P., Arendt, T., & Morawski, M. (2014). Aggrecan, link protein and tenascin-R are essential components of the perineuronal net to protect neurons against iron-induced oxidative stress. Cell Death Dis, 5(3), e1119. 10.1038/cddis.2014.25

Taylor, T., Dluhy, R. G., & Williams, G. H. (1983). β-Endorphin Suppresses Adrenocorticotropin and Cortisol Levels in Normal Human Subjects. The Journal of Clinical Endocrinology & Metabolism, 57(3), 592–596. 10.1210/jcem-57-3-592

Uchida, T., Furukawa, T., Iwata, S., Yanagawa, Y., & Fukuda, A. (2014). Selective loss of parvalbumin-positive GABAergic interneurons in the cerebral cortex of maternally stressed Gad1-heterozygous mouse offspring. Transl Psychiatry, 4(3), e371. 10.1038/tp.2014.13

van de Looij, Y., Larpin, C., Cabungcal, J. H., Sanches, E. F., Toulotte, A., Do, K. Q., & Sizonenko, S. V. (2019). Nutritional Intervention for Developmental Brain Damage: Effects of Lactoferrin Supplementation in Hypocaloric Induced Intrauterine Growth Restriction Rat Pups. Front Endocrinol (Lausanne), 10, 46. 10.3389/fendo.2019.00046

Vanhorebeek, I., Peeters, R. P., Vander Perre, S., Jans, I., Wouters, P. J., Skogstrand, K., Hansen, T. K., Bouillon, R., & Van den Berghe, G. (2006). Cortisol Response to Critical Illness: Effect of Intensive Insulin Therapy. The Journal of Clinical Endocrinology & Metabolism, 91(10), 3803–3813. 10.1210/jc.2005-2089

Vigetti, D., Deleonibus, S., Moretto, P., Karousou, E., Viola, M., Bartolini, B., Hascall, V. C., Tammi, M., De Luca, G., & Passi, A. (2012). Role of UDP-N-acetylglucosamine (GlcNAc) and O-GlcNAcylation of hyaluronan synthase 2 in the control of chondroitin sulfate and hyaluronan synthesis. J Biol Chem, 287(42), 35544–35555. 10.1074/jbc.M112.402347

Vizurraga, A., Adhikari, R., Yeung, J., Yu, M., & Tall, G. G. (2020). Mechanisms of adhesion G protein-coupled receptor activation. J Biol Chem, 295(41), 14065–14083. 10.1074/jbc.REV120.007423

Watanabe, H., Gao, L., Sugiyama, S., Doege, K., Kimata, K., & Yamada, Y. (1995). Mouse aggrecan, a large cartilage proteoglycan: protein sequence, gene structure and promoter sequence. Biochem J, 308 (Pt 2)(Pt 2), 433–440. 10.1042/bj3080433

Weber, P., Bartsch, U., Rasband, M. N., Czaniera, R., Lang, Y., Bluethmann, H., Margolis, R. U., Levinson, S. R., Shrager, P., Montag, D., & Schachner, M. (1999). Mice deficient for tenascin-R display alterations of the extracellular matrix and decreased axonal conduction velocities in the CNS. J Neurosci, 19(11), 4245–4262. 10.1523/jneurosci.19-11-04245.1999

Wieczorek, A., Perani, C. V., Nixon, M., Constancia, M., Sandovici, I., Zazara, D. E., Leone, G., Zhang, M. Z., Arck, P. C., & Solano, M. E. (2019). Sex-specific regulation of stress-induced fetal glucocorticoid surge by the mouse placenta. Am J Physiol Endocrinol Metab, 317(1), E109–e120. 10.1152/ajpendo.00551.2018

Willis, A., Pratt, J. A., & Morris, B. J. (2021). BDNF and JNK-signalling modulate cortical interneuron and perineuronal net development: implications for schizophrenia-linked 16p11.2 duplication syndrome. Schizophrenia Bulletin, 47, 812–816. 10.1093/schbul/sbaa139

Willis, A., Pratt, J. A., & Morris, B. J. (2022). Enzymatic Degradation of Cortical Perineuronal Nets Reverses GABAergic Interneuron Maturation. Mol Neurobiol, 59(5), 2874–2893. 10.1007/s12035-022-02772-z

Yu, Z., Han, Y., Hu, D., Chen, N., Zhang, Z., Chen, W., Xue, Y., Meng, S., Lu, L., Zhang, W., & Shi, J. (2022). Neurocan regulates vulnerability to stress and the anti-depressant effect of ketamine in adolescent rats. Molecular Psychiatry, 27(5), 2522–2532. 10.1038/s41380-022-01495-w

Zhang, S., Hu, S., Dong, W., Huang, S., Jiao, Z., Hu, Z., Dai, S., Yi, Y., Gong, X., Li, K., Wang, H., & Xu, D. (2023). Prenatal dexamethasone exposure induces anxiety- and depressive-like behavior of male offspring rats through intrauterine programming of the activation of NRG1-ErbB4 signaling in hippocampal PV interneurons. Cell Biol Toxicol, 39(3), 657–678. 10.1007/s10565-021-09621-0

Zhang, W., Watson, C. E., Liu, C., Williams, K. J., & Werth, V. P. (2000). Glucocorticoids induce a near-total suppression of hyaluronan synthase mRNA in dermal fibroblasts and in osteoblasts: a molecular mechanism contributing to organ atrophy. Biochem J, 349(Pt 1), 91–97. 10.1042/0264-6021:3490091

Zhang, Y., Chen, K., Sloan, S. A., Bennett, M. L., Scholze, A. R., O’Keeffe, S., Phatnani, H. P., Guarnieri, P., Caneda, C., Ruderisch, N., Deng, S., Liddelow, S. A., Zhang, C., Daneman, R., Maniatis, T., Barres, B. A., & Wu, J. Q. (2014). An RNA-sequencing transcriptome and splicing database of glia, neurons, and vascular cells of the cerebral cortex. J Neurosci, 34(36), 11929–11947. 10.1523/jneurosci.1860-14.2014

Zhong, Y., & Bellamkonda, R. V. (2007). Dexamethasone-coated neural probes elicit attenuated inflammatory response and neuronal loss compared to uncoated neural probes. Brain Res, 1148, 15–27. 10.1016/j.brainres.2007.02.024

Zhou, X. H., Brakebusch, C., Matthies, H., Oohashi, T., Hirsch, E., Moser, M., Krug, M., Seidenbecher, C. I., Boeckers, T. M., Rauch, U., Buettner, R., Gundelfinger, E. D., & Fässler, R. (2001). Neurocan is dispensable for brain development. Mol Cell Biol, 21(17), 5970–5978. 10.1128/mcb.21.17.5970-5978.2001

Zhu, B., Cui, G., Zhang, Q., Cheng, X., & Tang, S. (2019). Desumoylation of aggrecan and collagen II facilitates degradation via aggrecanases in IL-1β-mediated osteoarthritis. J Pain Res, 12, 2145–2153. 10.2147/jpr.S194306

Zhu, B., Luo, R., Jin, P., Li, T., Oak, H. C., Giera, S., Monk, K. R., Lak, P., Shoichet, B. K., & Piao, X. (2019). GAIN domain-mediated cleavage is required for activation of G protein-coupled receptor 56 (GPR56) by its natural ligands and a small-molecule agonist. J Biol Chem, 294(50), 19246–19254. 10.1074/jbc.RA119.008234

