## Supplementary information for "Glucocorticoids modulate expression of perineuronal net component genes and parvalbumin during development of mouse cortical neurons"

Supplementary Table 1

| Target gene | Forward primer sequences (5'-3') | Reverse primer sequences (5'-3') | Supplier |
|---|---|---|---|
| *Acan* | GTTGCAGACATTGACGAGTGC | AGTCCACCCCTCCTCACATT | Merck |
| *Bcan* | GATTCCGGGGTCTATCGCTG | ACGACCCCTTTGACCTTGAC | Merck |
| *Ncan* | AGTATGGGGGCCGGATCTGT | TGGTGTCCTGTGTGTCCTGAT | Merck |
| *Vcan* | ACCTTCCAACTATCCGGTGC | GGTATGCAGATGGGTTCATG | Merck |
| *Ptprz1* | TGCCGCCTGGATAAACCTCTC | CGGTGAAGTTGGGAAGCTGA | Merck |
| *Has1* | TGTGTCCTGCATCAGTGGTC | TTGGTGAGGTGCCTGTCATC | Merck |
| *Has2* | GCCATGTGGTTTCACAAGCA | TGAGACCCACTAGCTGGACA | Merck |
| *Has3* | GCTTCTTTGTGTGGCGTAGC | AGTCCACTGAGTTGCCAAGG | Merck |
| *Hapln4* | TCTGGAAGGGGTGGTCTTCC | AAAATGCCATCCTGTTCGGC | Merck |
| *TnR* | GGATATGCGGGATGGACAGG | GAGTCTCCTGCAGTGCCATT | Merck |
| *Pvalb* | CAAGCAGTCAGCGCCACTTA | GCGCAAAAGTCCTGTGTGTT | Merck |
| *Gad1* | TTTGGAGCTGTCTGACCACC | AAATCGAGGGTGACCTGTGC | Merck |
| *Gad2* | TCCTCTCTTGGCTGTAGCTGA | AGAGTTGGCCCTCTCTACTCC | Merck |
| *Gapdh* | AATGTGTCCGTCGTGGATCT | AGACAACCTGGTCCTCAGTG | Merck |
| *Tbp* | TGCTGTTGGTGATTGTTGGT | AACTGGCTTGTGTGGGAAAG | Merck |

Primer sequences for qPCR.

Supplementary Table 2

| Antibody | Species | Dilution | Company/supplier |
|---|---|---|---|
| *Bcan* | Mouse | 1: 1000 | Boehringer |
| *Has1* | Rabbit | 1: 1000 | Invitrogen |
| *Has2* | Mouse | 1: 5000 | Genetex |
| *Has3* | Rabbit | 1: 1000 | Genetex |
| *Hapln4* | Rabbit | 1: 1000 | St John's Laboratory |
| *TnR* | Rabbit | 1: 1000 | Genetex |
| *Gad65/67* | Rabbit | 1:10000 | Sigma-Aldrich |
| *Gapdh*-HRP conjugated | Rabbit | 1:10000 | Genetex |
| Anti-Mouse IgG HRP | Goat | 1:10000 | Cell Signaling |
| Anti-Rabbit IgG HRP | Goat | 1:6000 | Cell Signaling |

Dilutions and sources of antibodies used in western blot experiments

The efficiency of amplification of novel targets was calculated (Fig.1). Efficiency (%) of targeted primers ranged from 90% to 117%.

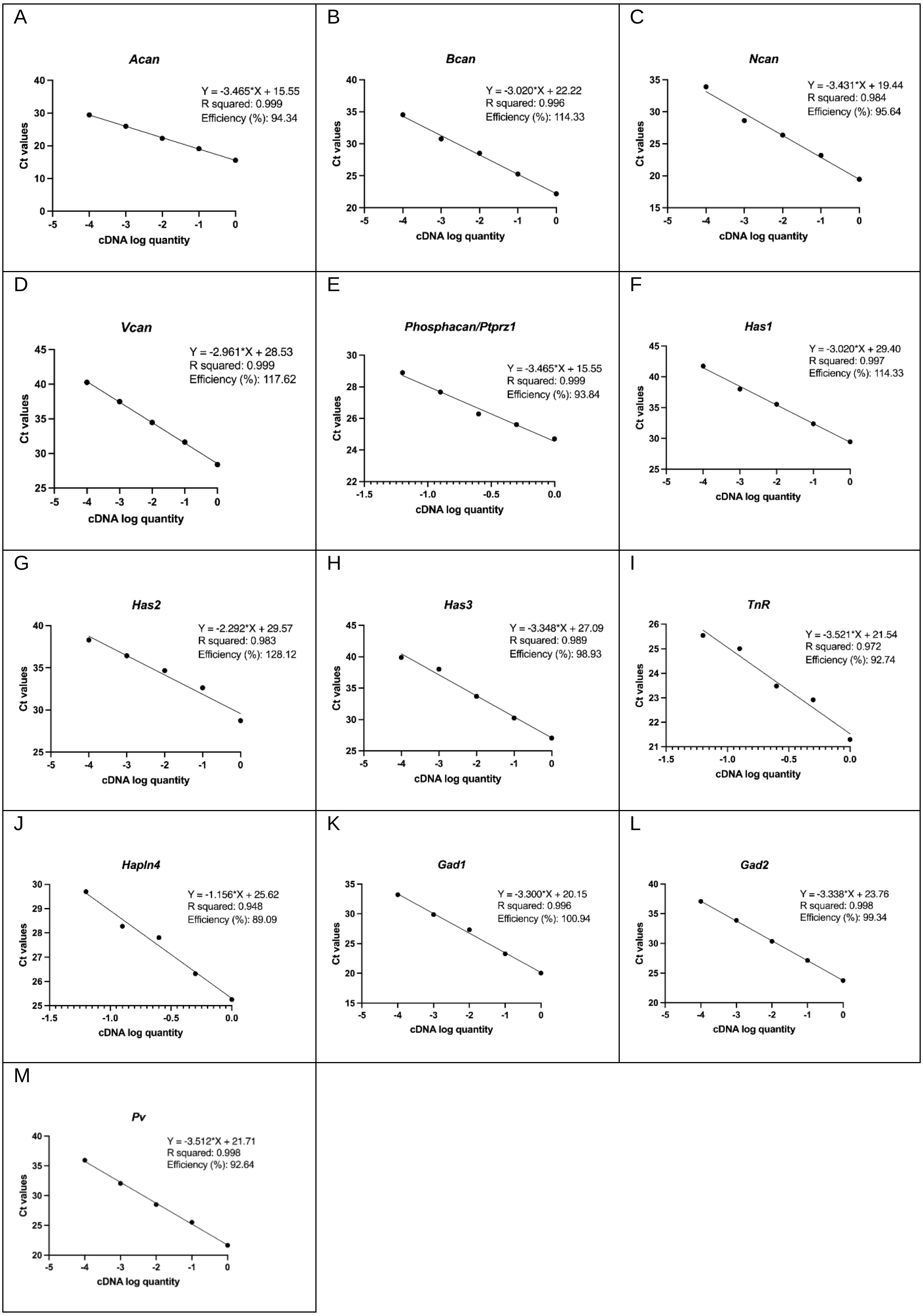


**Figure S1**: efficiency curves of targeted primers: A-J efficiency curves of PNNs-related primers; K-M: efficiency curves of GABA-related primers

Initial experiments established that *Gapdh* expression did not appear to be systematically affected by any of the treatments (Fig.S2), consistent with previous studies in the laboratory (e.g. Willis et al Schizophrenia Bulletin 2021).

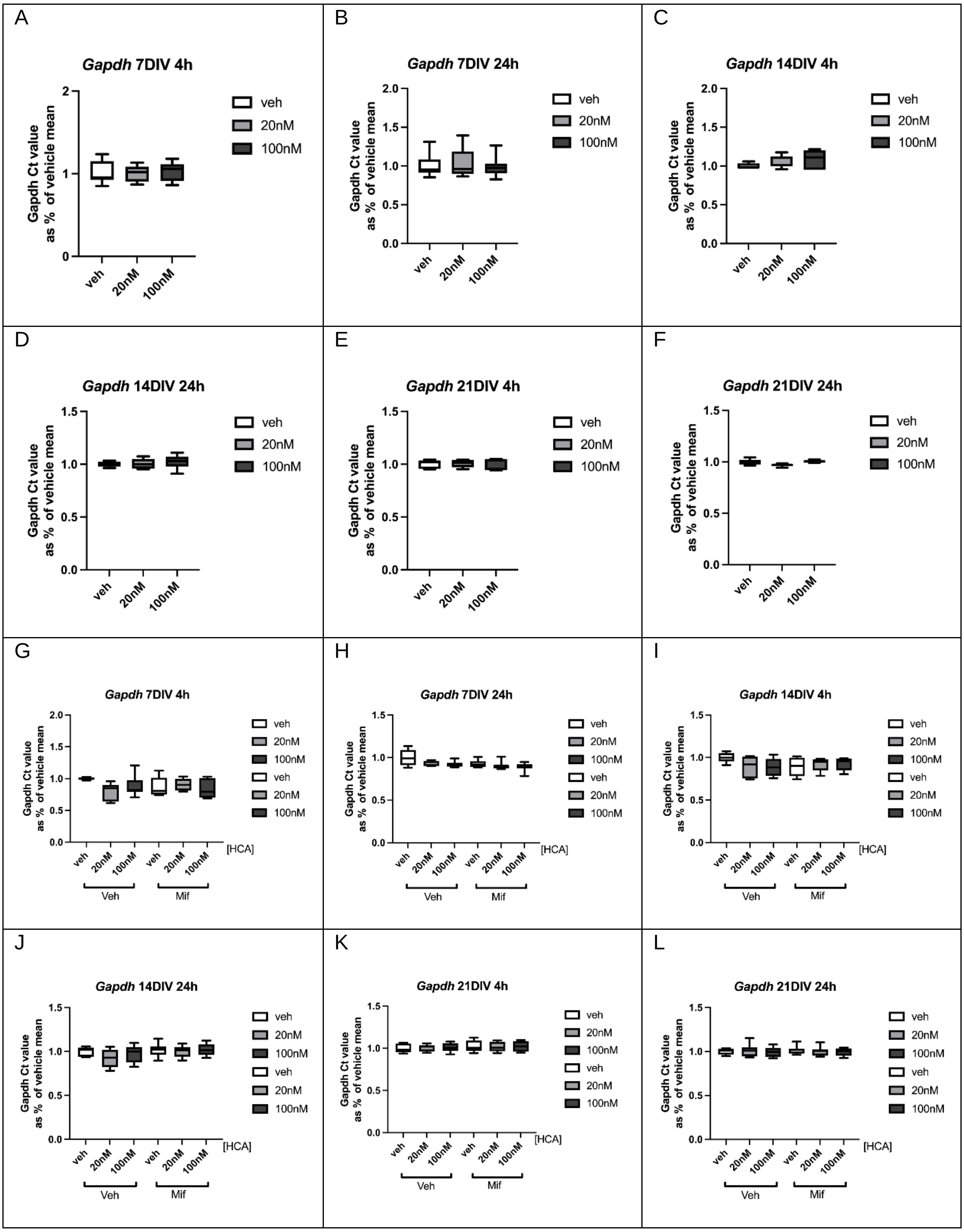


**Figure S2.** mRNA levels of *Gapdh* after exposure to low dose (20nM) and high dose (100nM) HCA for 4h and 24h. The mRNA levels remained unchanged at 7, 14 and 21 DIV.

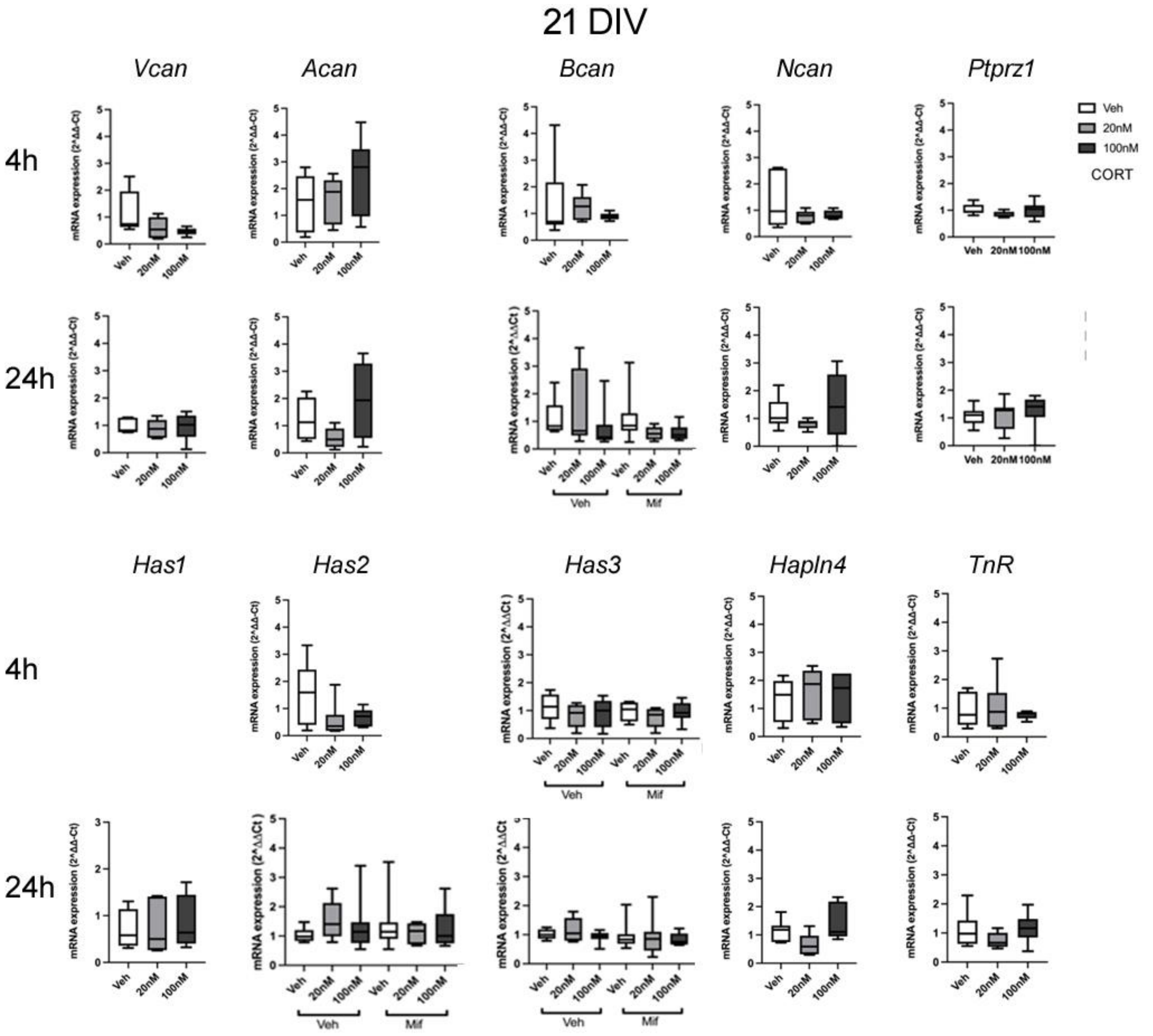


**Figure S3**: Lack of significant change in PNN component mRNAs when cortical cultures are treated with 20nM or 100nM CORT, or 20nM mifepristone, for 4h or 24h at 21 DIV. n=6-8/group. Boxes show median and interquartile range, with whiskers from minimum to maximum.

For *phosphacan* (*Ptprz1*), mRNA expression was also examined after 4h and 24h CORT exposure with low and high doses at 7 DIV, 14 DIV and 21 DIV. There were no significant changes of *Ptprz1* expression after CORT exposure (7DIV: 4h: $F$ (2, 43)=0.63 $p$=0.542, 24h: $F$ (2, 45) =1.78, $p$=0.66; 14 DIV: 4h: $F$ (2, 23)=1.21 $p$=0.39, 24h: $F$ (2, 19)=0.49 $p$=0.623; 21 DIV: 4h: $F$ (2, 18)=0.68 $p$=0.219, 24h: $F$ (2, 20)=0.55 $p$=0.584) (Figures S3, S4).

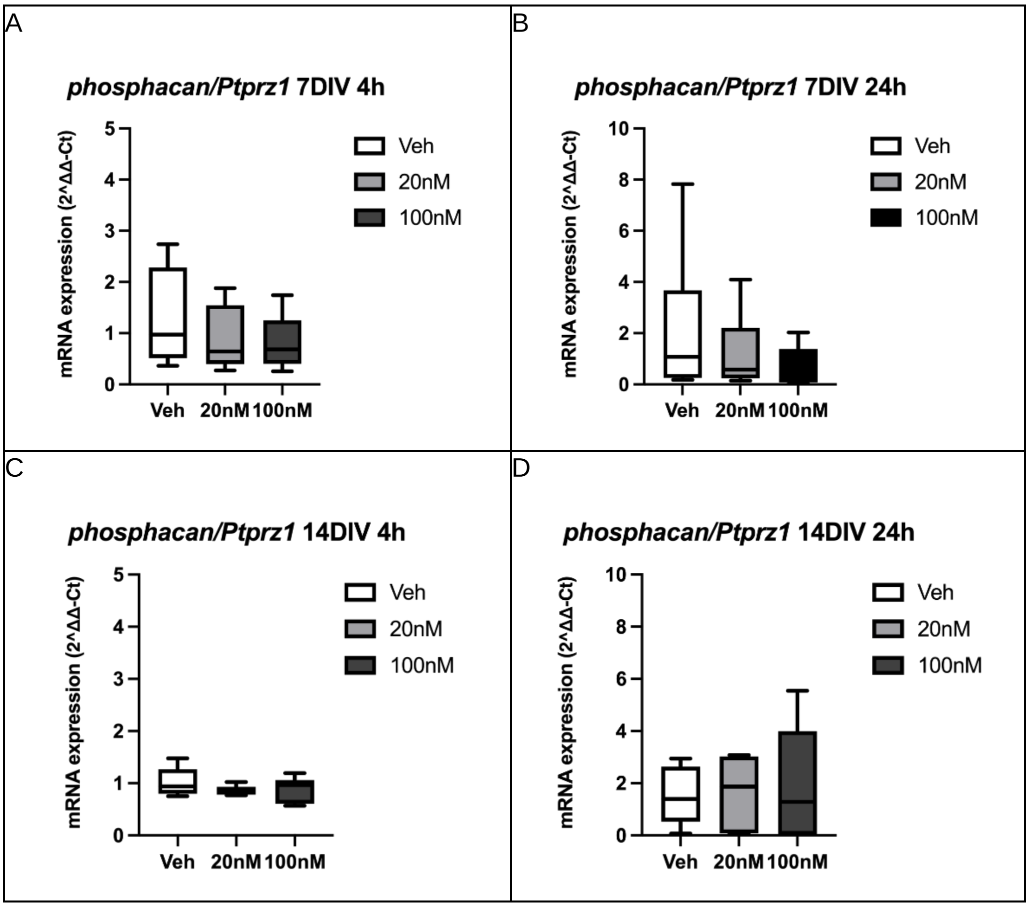


**Figure S4**: mRNA expression of *phosphacan/Ptprz1* after 4h and 24h exposure to low dose (20nM) and high dose (100nM) CORT at 7 and 14 DIV. There were no significant changes of PTPRZ expression detected at 7 DIV, 14 DIV and 21 DIV. (7 DIV: n=43 in total, veh=16, low dose=14, high dose=14; 14 DIV: n=20 in total, veh=6, low dose=8, high dose=7. Boxes show median and interquartile range, with whiskers from minimum to maximum.

With regard to *Hapln4* gene expression, no significant changes were detected at 7 DIV (4h: F (2, 11) =2.94, p=0.104, 24h: F (2, 8) =0.82, p=0.483) (Figure S5), or at 21 DIV (F (2, 18) =0.22, p=0.801) (Figure S3). However, the expression decreased significantly at 14 DIV (F (2, 15) =3.79, p=0.050, veh vs high dose CORT, p=0.048, Tukey post-hoc tests) 24h after high dose CORT treatment (not shown). Following mifepristone treatment at 14 DIV, the effect of CORT was also significant after 4h (F (2, 46) =5.91, p=0.006), but not 24h (F (2, 46) =0.18, p=0.836) (Figure S5). The mRNA expression of *Hapln4* decreased after low dose CORT treatment after 4h (veh vs low dose CORT p=0.026 Tukey post-hoc tests), this result was consistent with the previous findings. However, the overall effect of mifepristone was not significant at either 4h (F (2, 46) =0.14, p=0.714) or 24h (F (2, 46) =0.11, p=0.745). Similarly, the interactions of CORT and mifepristone were not significant either at 4h (F (2, 46) =0.69, p=0.506) or 24h (F (2, 46) =1.17, p=0.323). These results demonstrated that the effect of CORT was not mediated by the activation of GRs.

Based on the findings of mRNA alterations, protein levels of *Hapln4* were measured with CORT and mifepristone cotreatment at 14 DIV. The protein levels of *Hapln4* remained unchanged after 4h and 24h CORT exposure (4h F (2, 47) =0.23, p=0.794, 24h: F (2, 47)=0.80, p=0.455) (Fig.9 K L). Additionally, the overall effect of mifepristone (4h F (1, 47) =2.01, p=0.763, 24h: F (1, 47)=0.95, p=0.336) and the interactions of CORT and mifepristone (4h F(2, 47)=0.24, p=0.786, 24h: F(2, 47)=1.38, p=0.263) were not detected after 4h and 24h exposure, indicating the protein levels were not affected by GCs (Figure S5).

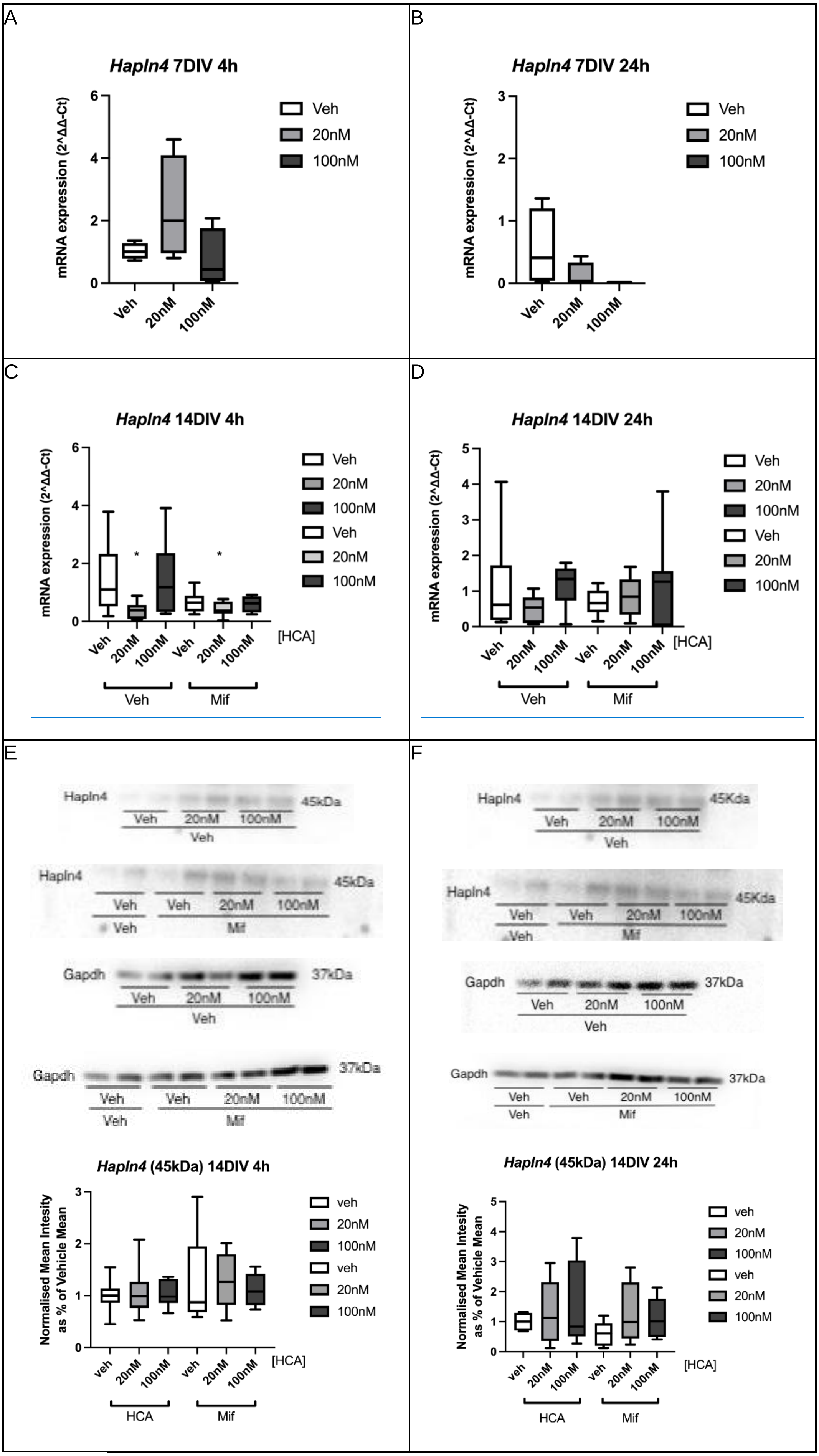


**Figure S5**: A-D: mRNA expression of *Hapln4* after 4h and 24h exposure to low dose (20nM) and high dose (100nM) CORT at 7, 14 and 21 DIV (7 DIV: n=20 in total, veh=8, low dose=6, high dose=7; 14 DIV: n=28 in total, veh=9, low dose=11, high dose=9; 21 DIV: n=39 in total, veh=14, low dose=14, high dose=12). G-J: mRNA expression of *Hapln4* at 14 and 21DIV after low dose (20nM) and high dose (100nM) CORT and mifepristone (100nM) treatment (n=48 in total, Veh: veh=8, low dose samples=8, high dose samples=8; Mifepristone: veh=8, low dose samples=8, high dose samples=8). E-F: Protein expression of *Hapln4* after low dose (20nM) and high dose (100nM) CORT and mifepristone exposure and relative box plots of normalised mean intensity of the bands' signals of *Hapln4*. (n=96 in total: veh: veh=16, low dose CORT=16, high dose CORT=16; Mifepristone: veh=16, low dose CORT=16, high dose CORT=16). *p<0.05 vs vehicle group, post-hoc Tukey's test. Boxes show median and interquartile range, with whiskers from minimum to maximum.

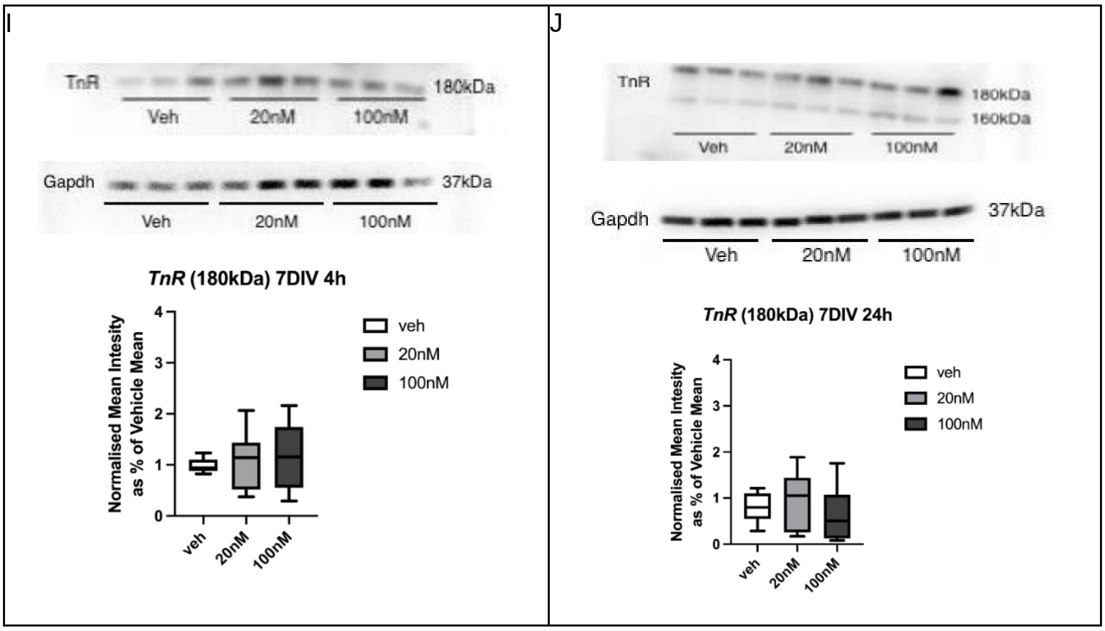


**Figure S6**: A-F: TnR protein expression after exposure to CORT. *TnR* 160KDa protein expression had an increasing tendency after CORT treatment. Western blot analysis of *TnR* proteins with CORT at 7 DIV for 4h and 24h respectively, with veh, low dose CORT (20nM) and high dose CORT (100nM), and relative box plots of normalised mean intensity of the bands signals of *TnR*. (n=27 in total, veh=9, low dose CORT=9, high dose CORT=9). K-L: Protein expression of *TnR* after low dose (20nM) and high dose (100nM) CORT and mifepristone (100nM) exposure, and relative box plots of normalised mean intensity of the bands signals of *TnR*. (n=96 in total: veh: veh=16, low dose CORT=16, high dose CORT=16; Mifepristone: veh=16, low dose CORT=16, high dose CORT=16). * p<0.05 vs corresponding vehicle group, post-hoc Tukey's test; # p<0.05, ## p<0.01 overall effect mifepristone vs vehicle group (ANOVA). Boxes show median and interquartile range, with whiskers from minimum to maximum.

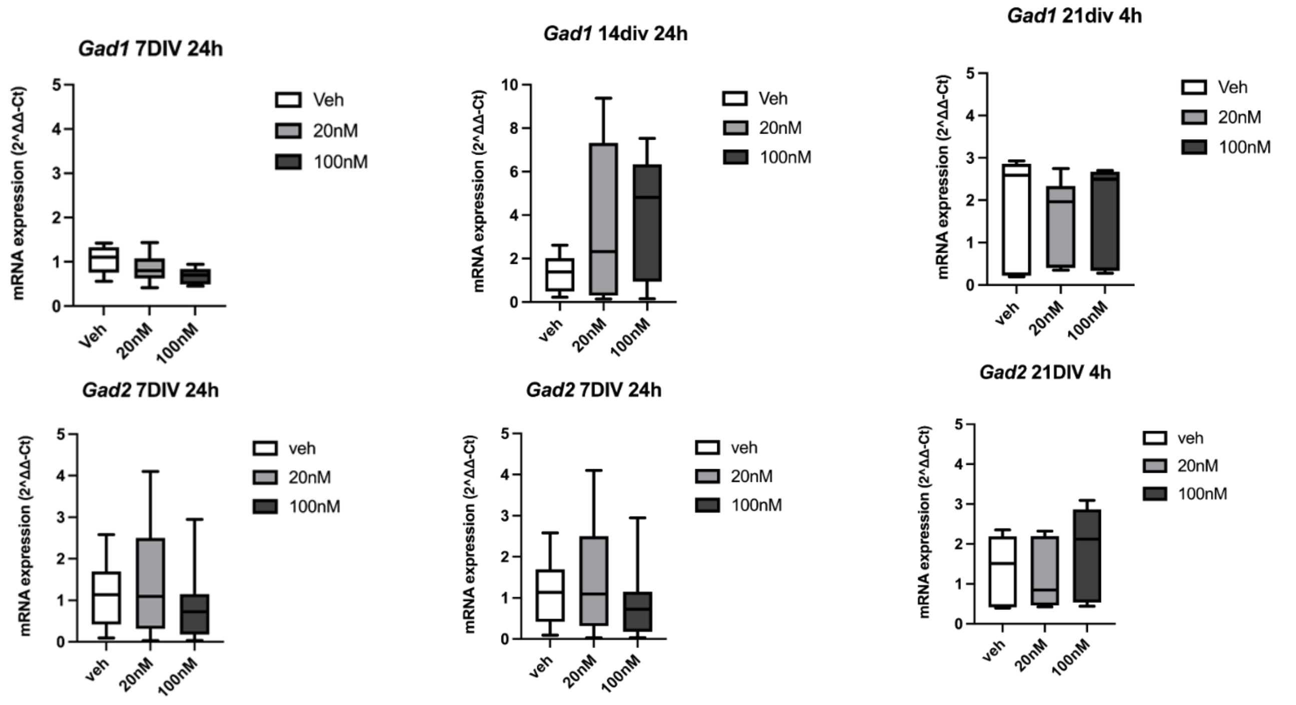


**Figure S7**: mRNA expression of *Gad1* (upper row) and *Gad2* (lower row), after treatment for 24h at 7DIV (left panels) and at 14 DIV (centre panels), and for 4h at 21 DIV (right panels) with low dose (20nM) or high dose (100nM) HCA at 7, 14 and 21 DIV. (7 DIV: n=20 in total, veh=8, low dose=6, high dose=7; 14 DIV: n=28 in total, veh=9, low dose=11, high dose=9; 21 DIV: n=39 in total, veh=14, low dose=14, high dose=12.). Boxes show median and interquartile range, with whiskers from minimum to maximum.

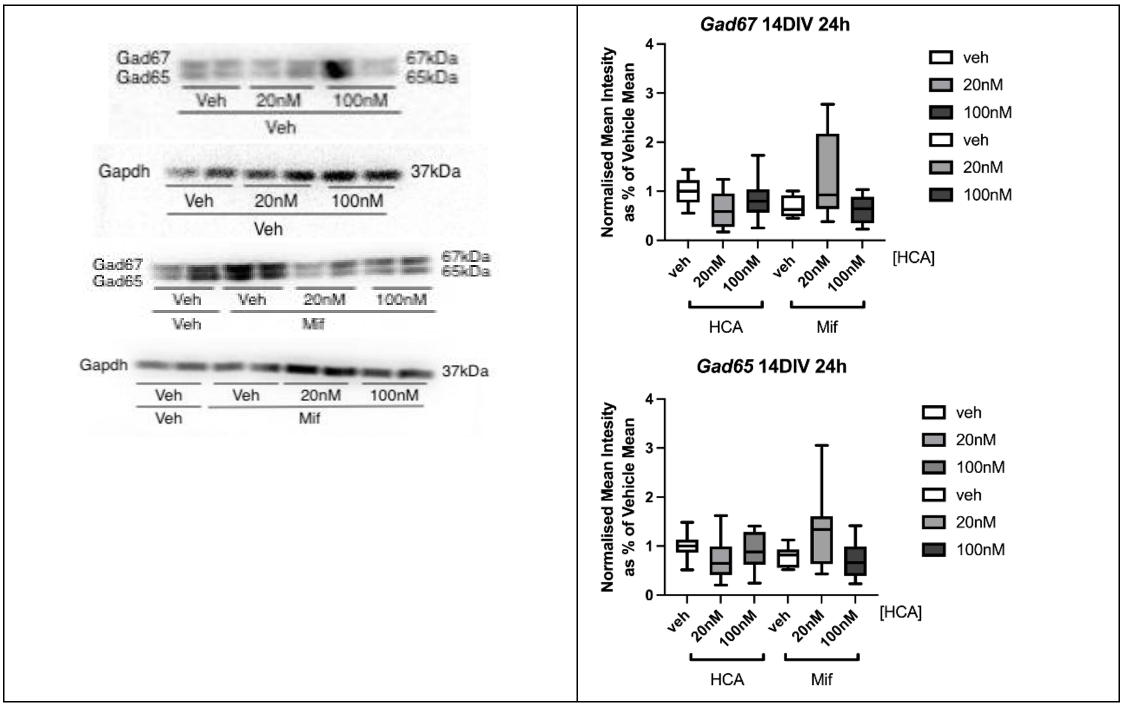


**Figure S8**: Gad65/67 protein levels at 14 DIV after 24h treatment. CORT had no effect on Gad1/Gad67 or Gad2/Gad65 after 24h exposure: Gad67: 24h F (2, 47) =0.68, p=0.512; Gad65: F(2, 47)=0.03, p=0.861). Mifepristone also had no effect on Gad1/Gad67 (F (1, 47) =0.03, p=0.861) or on Gad2/Gad65 (F(1, 47)=2.67, p=0.110) with 24h exposure. No interactions between CORT and mifepristone treatments was found for Gad1/Gad67 or Gad2/Gad65 (Gad67: F (2, 47)=1.50, p=0.235; Gad65: F(2, 47)=1.50, p=0.235). Boxes show median and interquartile range, with whiskers from minimum to maximum.

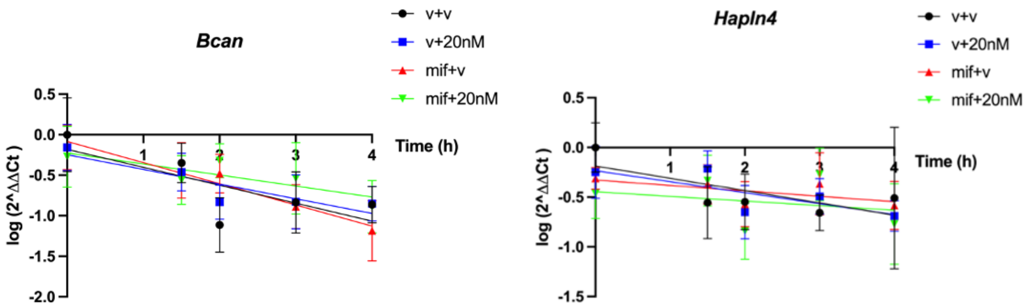


**Figure S9:** Measurement of mRNA decay with CORT (20nM) and mifepristone (20nM), for *Bcan* (left panel) and *Hapln4* (right panel). mRNA synthesis was inhibited by Act D (5µg/ml) for 0h, 1.5h, 2h, 3h and 4h. (Veh: veh: n=4, CORT: n=4; Mif: n=4, CORT: n=4). Mean +/- s.e.m. is shown.

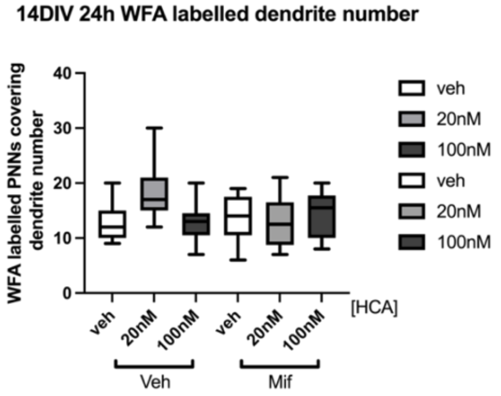


**Figure S10**: Lack or effect of CORT or mifepristone on the number of neuronal dendrites per cell covered by PNNs labelled by WFA at 14DIV. Graph shows the number of dendrites/cell covered by PNNs, quantified after 24h treatment with veh+veh (distilled water), veh+20nM CORT and veh+100nM CORT, and Mif+veh, Mif+20nM CORT and Mif+100nM CORT. (Veh: veh: 139 dendrites in 13 cells nested in 3 different slides with 2 cultures, 20nM: 275 dendrites in 17 cells nested in 3 different slides with 2 cultures, 100nM: 164 dendrites in 15 cells nested in 3 different slides with 2 cultures; Mif: 290 dendrites in 21 cells nested in 3 different slides with 2 cultures, 20nM: 182 dendrites in 16 cells nested in 3 different slides with 2 cultures, 100nM: 177 dendrites in 14 cells nested in 3 different slides with 2 cultures). Boxes show median and interquartile range, with whiskers from minimum to maximum.
